# Ramp-shaped neural tuning supports graded population-level representation of the object-to-scene continuum

**DOI:** 10.1101/2022.01.06.475244

**Authors:** Jeongho Park, Emilie Josephs, Talia Konkle

## Abstract

We can easily perceive the spatial scale depicted in a picture, regardless of whether it is a small space (e.g., a close-up view of a chair) or a much larger space (e.g., an entire class room). How does the human visual system encode this continuous dimension? Here, we investigated the underlying neural coding of depicted spatial scale, by examining the voxel tuning and topographic organization of brain responses. We created naturalistic yet carefully-controlled stimuli by constructing virtual indoor environments, and rendered a series of snapshots to smoothly sample between a close-up view of the central object and far-scale view of the full environment (object-to-scene continuum). Human brain responses were measured to each position using functional magnetic resonance imaging. We did not find evidence for a smooth topographic mapping for the object-to-scene continuum on the cortex. Instead, we observed large swaths of cortex with opposing ramp-shaped profiles, with highest responses to one end of the object-to-scene continuum or the other, and a small region showing a weak tuning to intermediate scale views. Importantly, when we considered the multi-voxel patterns of the entire ventral occipito-temporal cortex, we found smooth and linear representation of the object-to-scene continuum. Thus, our results together suggest that depicted spatial scale is coded parametrically in large-scale population codes across the entire ventral occipito-temporal cortex.

## INTRODUCTION

When we see a picture of an environment, we can easily perceive the spatial scale depicted in the image. Even in images that are the same 2-dimensional size, some may depict a very small space (e.g., a close-up view of a chair; typically referred to as “objects” stimuli), whereas others may depict a much larger space (e.g., a view of an entire classroom; typically referred to as”scenes” stimuli). Although objects and scenes have been typically treated as contrasting categories with a clear distinction (Henderson and Hollingworth, 1999; Logothetis and Sheinberg, 1996; Oliva, 2005; Steeves et al., 2004; Epstein and Kanwisher, 1998; Grill-Spector et al., 2001), this example illustrates that they can also be considered as the opposite ends of a continuous dimension of depicted spatial scale (Henderson and Hollingworth, 1999). This continuum of views ranges from object to scene views, with numerous intermediate-scale views between the two extremes (hereinafter, the “object-to-scene continuum”). Here, we examined the tuning and topographic organization of brain responses along the visual system to views varying along this continuum, to gain insight into the underlying neural coding of depicted spatial scale.

There are well-established brain regions that show selective preference for either end of the object-scene continuum (for reviews see Epstein and Baker, 2019; Dilks et al., 2021; Gauthier and Tarr, 2016; Tacchetti et al., 2018). An object-preferring lateral occipital complex (LOC; Malach et al., 1995) is located on the lateral bank of the fusiform gyrus, and a scene-preferring parahippocampal place area (PPA; Epstein and Kanwisher, 1998) is located on the medial side of the inferior temporo-occipital cortex. A few studies have explored the response properties of these regions along different depicted scales (e.g. Park et al., 2015; Lescroart and Gallant, 2019; Persichetti and Dilks, 2016; Quinlan and Culham, 2007; Troiani et al., 2014)–e.g. Park et al., 2015 found slight but systematically stronger overall responses in scene-selective regions as the depicted scale increased from the size of a walk-in closet to a large-scale arena. And, in recent work considering brain responses evoked by intermediate “reachable-scale” views, we found regions that were activated more by images of reachable-scale environments (e.g., office desktops or kitchen counters) than either close-scale object views or navigable-scale scene views (Josephs and Konkle, 2020). This work also showed that object- and scene preferring regions both showed intermediate level of activation to reachspace views (compared to object and scene views). Taken together, current research has demonstrated that different parts of the ventral visual stream are sensitive to different parts of the object-to-scene continuum.

More generally, these findings raise the intriguing possibility that depicted spatial scale is encoded smoothly along the cortical surface, with adjacent cortical regions tuned to adjacent points along the continuum. That is, using a “multi-channel model”, the full object-to-scene continuum might be encoded through a bank of narrow Gaussian tuning functions, which show the highest activation to a particular depicted spatial scale, and increasingly less activation to views with a depicted spatial scales on either side of this peak. This kind of encoding scheme is evident in the representation orientation and spatial frequency in early visual cortex (Blakemore and Sutton, 1969; Pond et al., 2013). In the case of these feature dimensions, the spatial arrangement of neurons with different preferred orientations forms a radial-shape preference map (“orientation pinwheels”; Yacoub et al., 2008). Thus, one possibility for the tuning and topography of cortical responses to depicted spatial scale is that there is a continuous bank of response peaks to different points along the object-to-scene continuum, connecting the object-selective LOC, scene-selective PPA, and the more posterior ventral reachspace patch (vRSP) in between, in to one larger-scale map (Op De Beeck et al., 2008).

However, there are a number of other possible encoding schemes to represent continuous dimensions that do not rely on a multi-channel model. For example, depicted spatial scale may also be encoded more parametrically within a region, with monotonic tuning curves that could be either linear (i.e., ramp-shaped), or non-linear (e.g., sigmoidal). This kind of ramp-shaped coding has been hypothesized to underlie facial identity coding, where different sub-populations of neurons have monotonic neural response to the geometry of facial features, such as eye height or inter-eye distance (Susilo et al., 2010; Freiwald et al., 2009). Under this scheme, spatial scale could be captured at a population level with opposing populations, in which neural sub-populations respond maximally to either extreme of the dimension, the balance of which signals the depicted spatial scale. There are many hypothesized advantages of such opponent coding, including the possibility that it can be used to amplify behaviorally relevant category boundaries through winner-take-all mechanisms (Suzuki et al., 2005, p.161). This scheme would predict a more abrupt transition in tuning preferences along the cortical surface, as adjacent populations transition from near-preference to far-preference. However, this opponent coding model does not necessarily require or predict the existence of the intermediate-scale tuned regions (Josephs and Konkle, 2020). More generally, there are other theoretically possible kinds of response profiles as depicted scale varies (e.g. a region with bimodal tuning, with highest responses to both extremes). At stake with these different encoding schemes is how information about the depicted scale is formatted along the visual system, with corresponding implications for how it can be read-out from the structure brain responses.

To explore these possibilities, here we measured human brain responses using functional magnetic resonance imaging, while participants viewed a continuum of spatial scales. To systematically manipulate the spatial scale while controlling other potential confounding factors, we constructed virtual 3-dimensional naturalistic indoor environments with identical physical dimensions. Within each environment, we captured a smooth continuum of views by taking snapshots of the environment from a camera that moved from a far to a close distance relative to a center object. We then (i) characterized the response properties of the ventral stream along this object-to-scene continuum, (ii) differentiated the underlying tuning profiles using a data-driven clustering method, and (iii) explored the nature of the population code using simulations. To anticipate, we did not find evidence for a smooth topographic mapping of this continuum along the cortical surface, in which adjacent depicted scales have tight tuning functions along adjacent regions of cortex. Instead, we revealed two large adjoining zones of cortex showing the opposite ramp-shaped tunings, with a small posterior region with intermediate scale preferences. We discuss the implications of these findings, in which information about the spatial scale is implicit in the balance of activation among two opposing sub-populations, and we address the implications of this interpretation for the smaller regions which show preferring intermediate-scale views.

## RESULTS

### Cortical mapping of the object-to-scene continuum and underlying tuning profiles

Is there a large-scale topographic organization that smoothly maps the object-to-scene continuum (Figure 1A)? To investigate this question, we showed participants snapshots of virtual indoor environments that were captured at various viewing distances from a central object. We created 20 different environments that had identical physical size and spatial dimensions. The environments included several semantic scene categories (e.g., kitchens, bedrooms, offices, etc), and each environment was populated with both small (e.g., tools) and big objects (e.g., cabinets) that are consistent with its category. Importantly, by the back wall of each environment, there was a “central object” on top of some desk-like structure (e.g., Figure 1A).

**Figure 1:**
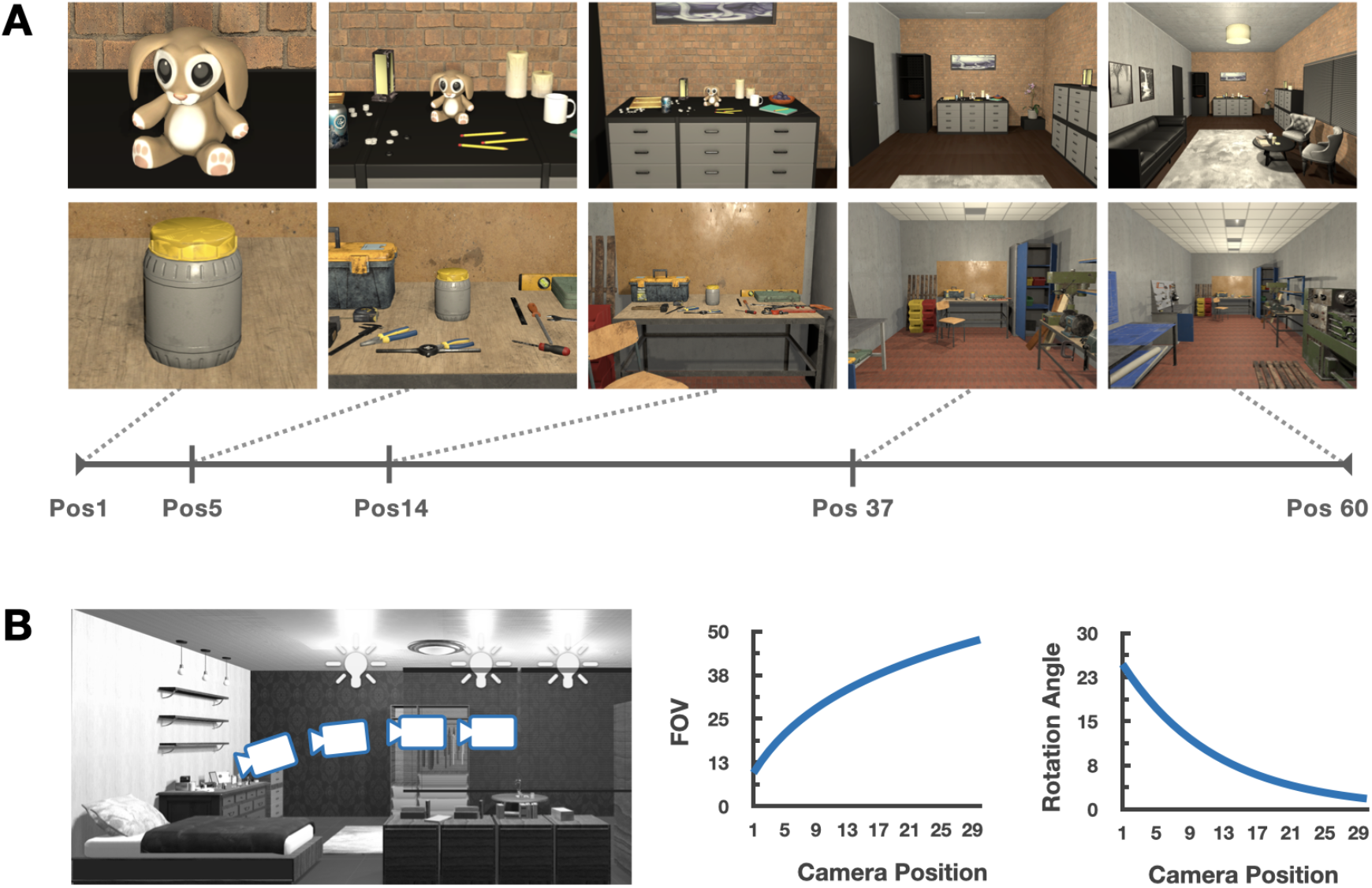
Stimuli. (A) Example images taken from two different environments (each row). Within each environment, we varied the the camera position along 15 log-scaled points, from a close-up object view (Position 1) up to a full-scene view (Position 60). Here, 5 positions are shown for illustration purposes. (B) Two camera parameters were systematically varied with the change of camera position. First, the field of view (FOV) was smallest at the closest view, then increased logarithmically as the camera moved toward the farthest view. Second, the rotation angle was downward at the closest view, then gradually adjusted upward, simulating a person looking at the central object from different positions in the room.

The viewing distance for a given snapshot was defined as the distance from this central object to the position of the camera. This viewing distance was used as a proxy for the depicted spatial scale of a view, and was the key manipulated variable in our experiments. We varied the position of camera along 15 log-scaled points spanning from directly in front of the object to the back of the room (see Figure 1B, left for a depiction of a sample views along this continuum). To ensure that the closest view was more object-focused, while the farthest scale view was more scene-focused, we had to make two further parallel adjustments to the camera parameters along this continuum. First, we adjusted the field of view of the camera, so that the closest view was a closer crop of the object, consistent with how object images are typically depicted to study object responses in the scanner; the field of view increased logarithmically with each step away from the object, to arrive at a more typical scene-focused view at the farthest position (Figure 1B, middle). Second, we also adjusted the angle of the camera as a function of position (simulating a person with a fixed height, who must angle their head down on the approach to an object); the camera was angled downward to center the object at the closest distance, and the angle was gradually adjusted upward until it was parallel to the floor plane (Figure 1B, right). See https://osf.io/hcmgk/ for all stimuli.

In the fMRI scanner, participants viewed images presented in a standard blocked design, while performing a one-back repetition detection task. A given block contained views from the same viewing distance (i.e., depicted spatial scale), sampled across different environments. Beta weights were extracted for each level of depicted spatial scale, for each voxel in each participant (Talairach space). Then, we computed the group-level betas by averaging betas across participants for each condition. Voxels used in subsequent analyses were selected based on the voxel-wise reliability within an anatomical mask (Tarhan and Konkle, 2020a), which included occipito-temporal cortex (OTC), occipito-parietal cortex, and the corresponding medial part of the brain (see Supplemental Figure 1A).

We first mapped the peak response preference of each voxel. Figure 2 shows these maps, in which each voxel was colored based on its preferred spatial scale (i.e., scale with the highest activation), where larger activation differences between the highest and the second highest condition are indicated with more saturated colors. The resulting preference map revealed extensive regions of cortex with a clear preference for either the closest object views or the farthest scene views. On the lateral surface, there was a large territory of cortex preferring the single-object view (dark red, Figure 2); on the medial side, there was another large territory preferring the far-scene view (dark blue). Although somewhat smaller, there was also some portion of cortex preferring the intermediate scale (yellow to green colors, Figure 2), in between the object-preferring and scene-preferring regions. While individual subject maps showed large individual differences (Supplemental Figure 2A), the two clusters preferring the extreme close object view along the lateral cortex, and the extreme far scene-view along the medial cortex were consistently observed (Supplemental Figure 2B). Overall, this preference map shows clear evidence for the preference of either extreme spatial scale, and some for the intermediate; we did not find clear evidence for a continuous mapping of the object-to-scene continuum across the cortical surface.

**Figure 2:**
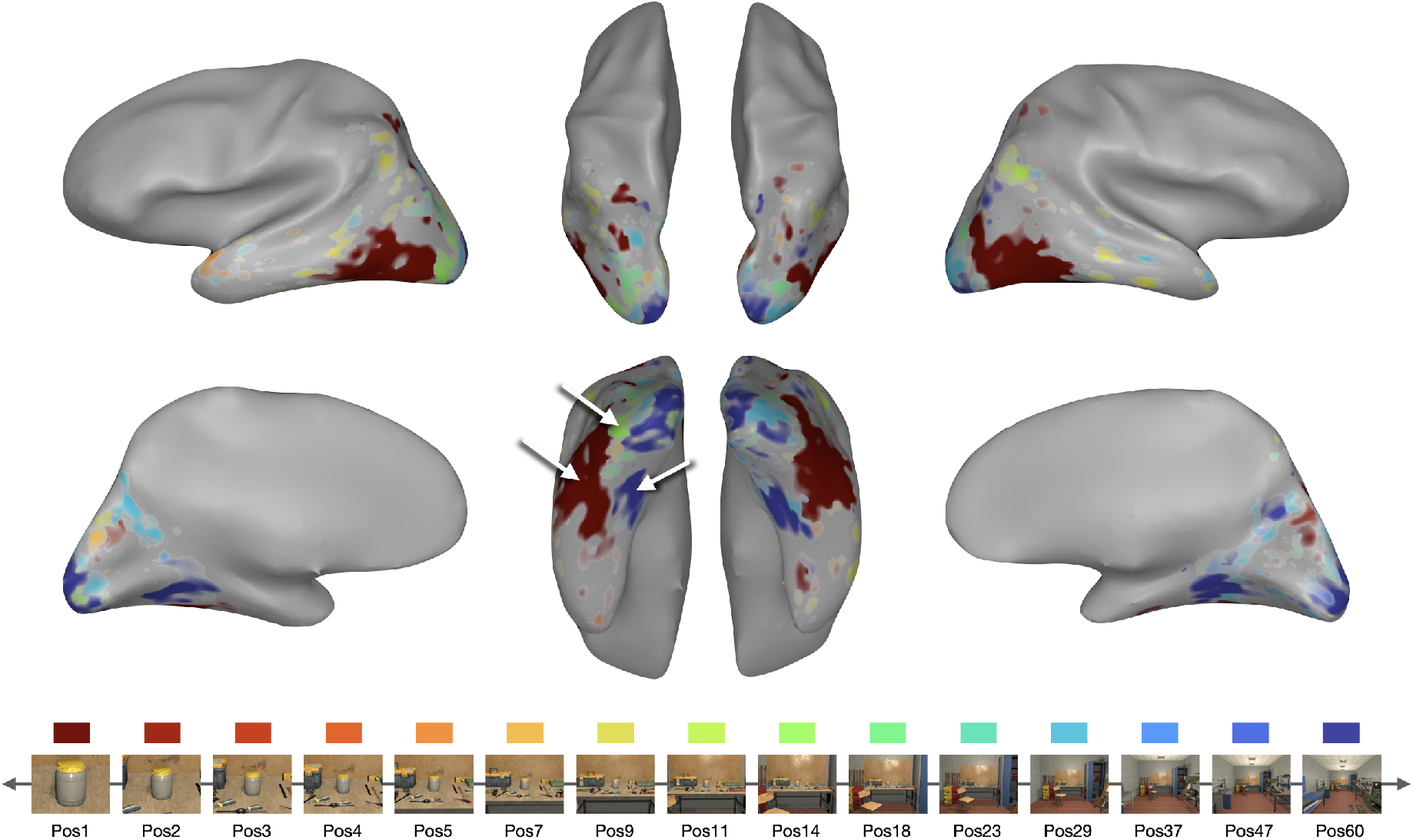
Preference map. Along the bottom, we show example images from 15 conditions depicting the object-to-scene continuum, and the corresponding color scheme for the plot above. The preference mapping result is visualized on an example subject brain, where each voxel is colored based on its most preferred condition (i.e., the highest activation), and the saturation is determined by the activation difference between the highest and the next highest condition. In the ventral occiptotemporal cortex (bottom middle), there is a large swath of cortex preferring the single-object views (dark red) and another large territory preferring the far-scene views (dark blue). There is also a small cluster preferring the intermediate-scale views (green).

The preference mapping approach groups voxels based on their highest activation, implicitly assuming a voxel tuning shape with a single-peak. This approach might obscure other theoretically interesting tuning profiles. For example, some voxels might have preference for two different conditions (e.g., bimodal tuning), or some might have a parametric tuning across the conditions (e.g., Park et al., 2015). To test various response profiles of voxels without making a priori assumptions about them, we used a data-driven clustering approach, called response profile clustering (RPC; Tarhan and Konkle, 2020b). Specifically, we grouped voxels that have similar response profile using *k*-means clustering. As there were no constraints to group voxels into contiguous clusters, this approach also allows for the natural organizational structure to be revealed, based on differences in tuning along the object-to-scene continuum.

Here we report the response profile clustering solution with four groups of voxels (*k*=4) and their corresponding response profiles. To select this number, we first computed a range of *k*-means solutions varying the number of clusters (*k*) from 2 to 20. We chose the final *k* based on the cluster center similarity (Supplemental Figure 3A), which measures how similar the cluster centers are to one another on average. To visualize the clustering solution (*k*=4), we created a cortical map, in which all voxels assigned to the same cluster were colored the same (Figure 3A) and the corresponding cluster centroids are plotted underneath (Figure 3B; Supplemental Figure 4 shows the results for a range of *k*s).

**Figure 3:**
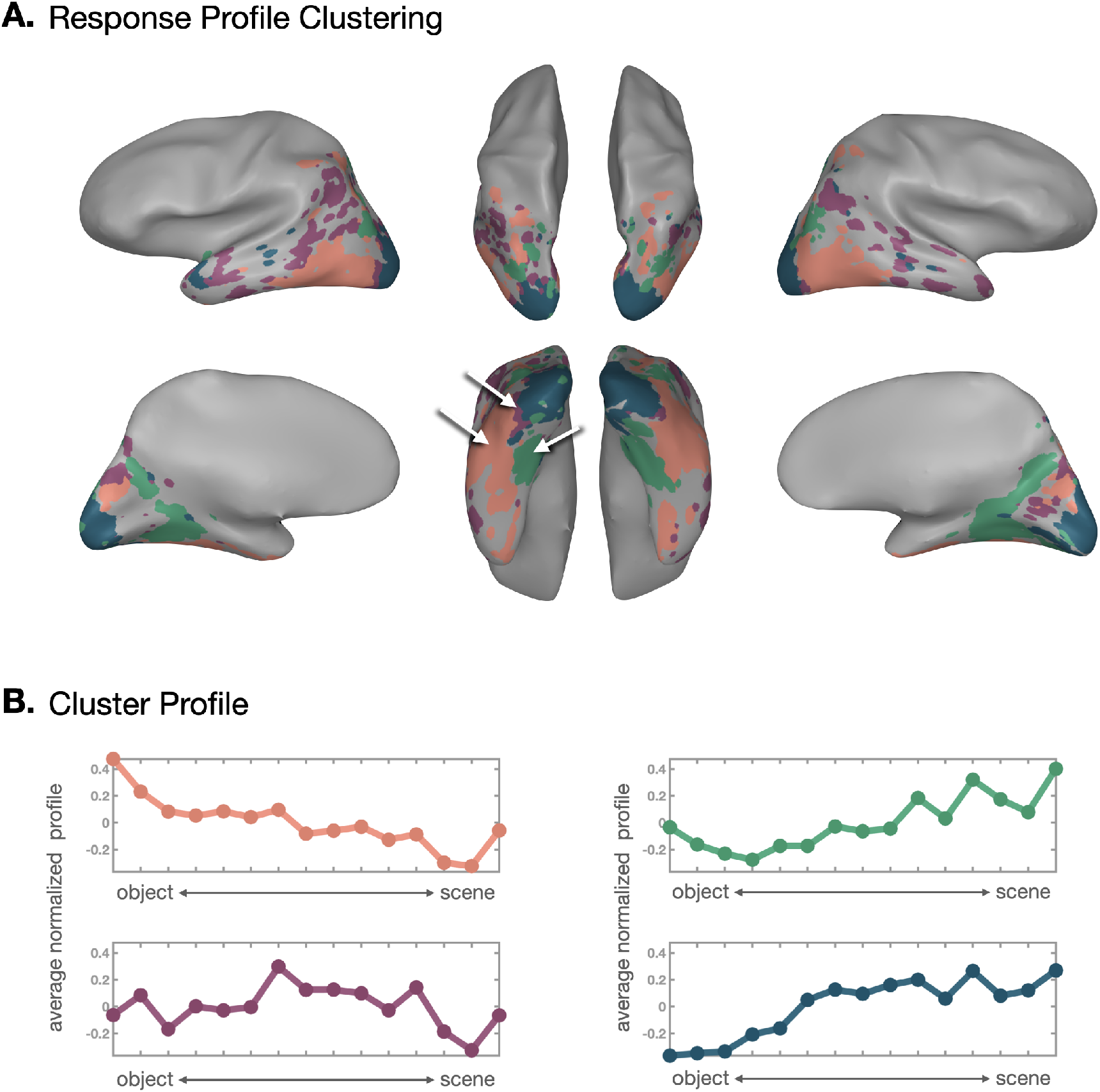
Response Profile Clustering. (A) The RPC results are visualized on an example subject brain (k=4). Each voxel is colored based on its assigned cluster, and clusters with more similar response profiles have colors that are more similar in hue. (B) The response profile corresponding to cluster centroids are plotted with the corresponding cluster color on the brain plot. The pink cluster shows the highest activation to single-object views and gradually less response toward the far-scene views. The green cluster shows the opposite pattern, where the highest activation was at the far-scene views. The purple cluster activation peaks for intermediate-scale views. The dark green cluster shows a similar but distinguishable response pattern from the green cluster, and it corresponds to early visual areas that were defined with independent localizer data.

**Figure 4:**
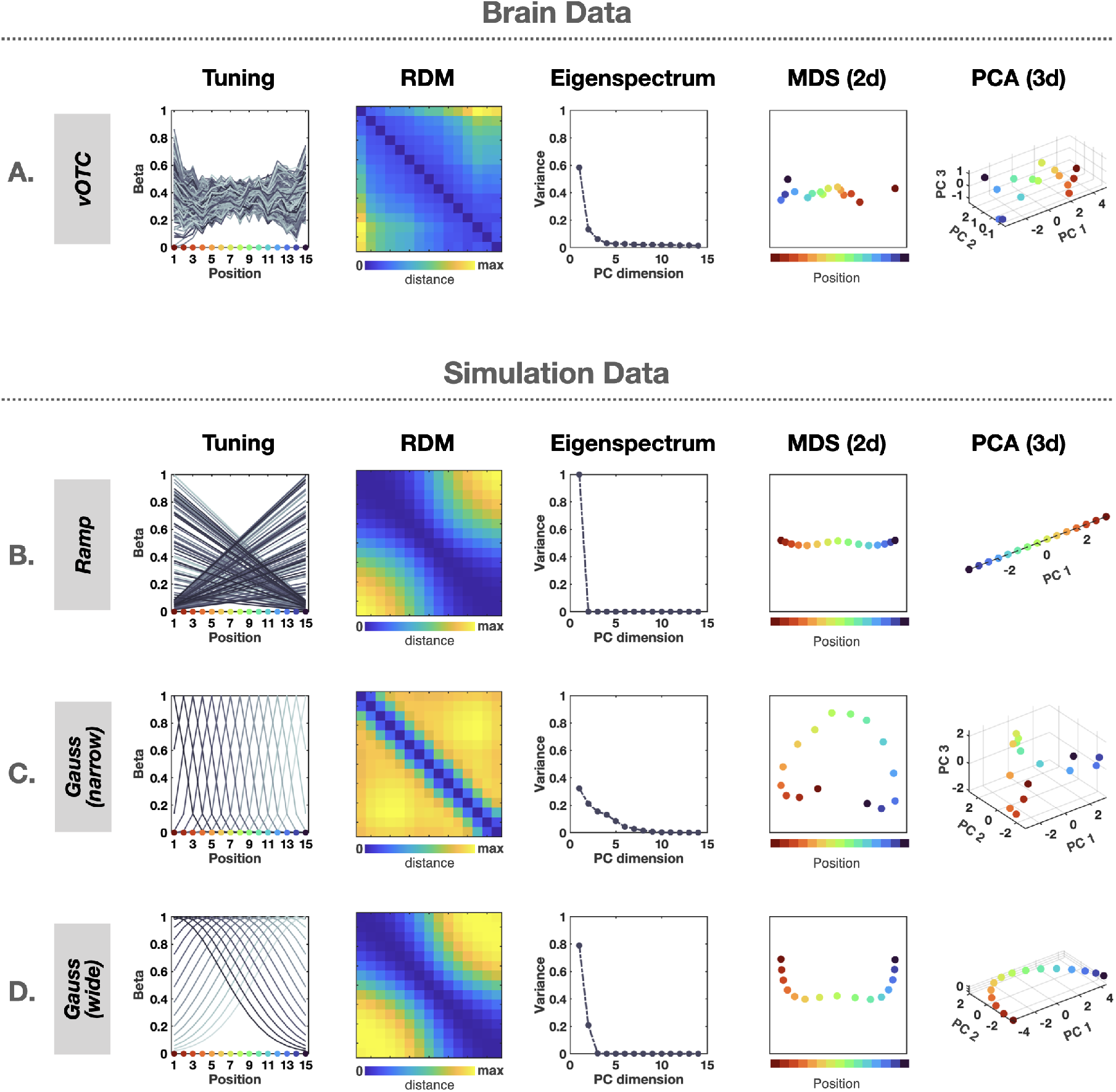
Neural tuning and representational geometry analysis. (A) Population of responses in the brain data. We show normalized voxel tunings over the object-to-scene continuum (Tuning), pairwise representational distances between conditions (RDM), how much variance is explained by each principal component (eigenspectrum), and visualization of representational geometry in the population (MDS (2d) and PCA (3d)). (B-D) Simulation Idealized voxel tunings and representational geometry. The ramp-shaped tuning (B) shows one-dimension feature space with a perfectly linear trajectory, whereas the narrow Gaussian tuning (C) shows much higher feature dimension with a cardioid-like trajectory. However, the wide Gaussian tuning (D) showed similar representational geometry to the ramp-shaped tuning.

Interestingly, three of the four data-driven clusters showed strong concordance with the zones which emerged from the preference mapping results. The first group of voxels (pink cluster) were mostly positioned at the lateral side of ventral occipital cortex. They formed a large-scale, spatially contiguous cluster, even though the clustering algorithm does not consider voxels’ anatomical locations or their spatial proximity to each other. This group of voxels showed the highest response to objects and gradually smaller response for scenes. The response change across the continuum was fairly smooth, resembling ramp-shaped tuning (c.f. Freiwald et al., 2009). In contrast, the second group of voxels (green cluster) covered a large swath of cortex at the medial side of ventral occipital cortex. This cluster showed the highest response to scenes and a graded response toward the object, with opposing ramp-shaped tuning. It is also noticeable that this cluster extended beyond the ventral region to the medial part where the retrosplenial cortex (RSC) is located, supporting this cluster’s preference for scenes. Results for additional category-selective regions are shown in Supplemental Figure 6. The third group of voxels (purple) is spatially positioned in between the first and second clusters, and showed the strongest response for intermediate-scale views. These three clusters were repeatedly found regardless of the number of clustering solutions (*k*) we choose, whereas other clusters were less robust and more variable depending on the *k*. The fourth group of voxels (dark green) near the occipital pole showed a similar but distinctive pattern from the green cluster. Crucially, this cluster corresponded to an early visual cortex (V1-V3), which was separately defined with an independent data.

How robust is this clustering solution? We quantified robustness of the results in two ways. First, we assessed the reliability of the clustering solution across different subsets of stimuli. Given *k*, the clustering was performed separately for two subsets of data which were divided by the stimuli (e.g., environment set A vs. environment set B). Then, the agreement of clustering solutions was quantified using the *d*-prime (see Methods). This analysis showed consistent clustering results across different environments (*d’* = 1.2, Supplemental Figure 3C), suggesting that the observed solution likely reflects some common features across environments (e.g., spatial scale or spatial layout) rather than environment-specific features (e.g., particular objects or textures). Second, we assessed the stability of the clustering solutions across participants. For each iteration (50 iterations), the participants were randomly split into two groups, and the sensitivity of one group to predict the other was computed using the *d’* (see Methods). The results showed moderate *d’* scores across the whole range of *k*s (0.86-0.98, mean = 0.88, s.d. = 0.03), indicating that all of the clustering solutions observed here are relatively stable (Supplemental Figure 3B).

Altogether, we did not find a smooth topographic mapping of the object-to-scene continuum in the peak responses along the cortical surface, considering both our preference mapping analysis and the more nuanced response profile clustering (RPC). Instead, these analyses revealed clear evidence for distinct swaths of cortical territory with opposing ramp-shaped tuning, with peak responses at either extreme of the object-to-scene continuum, and smaller clusters of voxels intermediate in anatomical position with intermediate tuning preferences. Given this kind of structure in the univariate response profiles of these voxels, we next examined the impact of such tunings on the representation of the object-to-scene continuum at *population level*–that is, how are the distinctions between depicted spatial scale realized in the distributed population responses that span these differently-tuned clusters.

### Population-level Response Structure

To investigate how depicted spatial scale is encoded in the population of the entire ventral occipital temporal cortex (vOTC) voxels, we conducted several analyses to characterize and visualize the representational geometry (Kriegeskorte and Wei, 2021), depicted in Figure 4A. First, a representational dissimilarity matrix (RDM) was generated by computing the correlation between multi-voxel patterns of each condition pair (Figure 4A). There was a smooth pattern along the diagonal in the RDM, indicating that images with similar spatial scales also have more similar neural patterns across vOTC voxels.

Next, inspired by analyses in Stringer et al., 2019, we conducted a principal component analysis (PCA) over the condition × voxel matrix, which gives a sense of the dimensionality of the population code over these conditions. The eigenspectrum—that is, the proportion of explained variance by each principal component—is shown in Figure 4A. In the population of vOTC voxels, the first PC explained 58.5% of variance, followed by the second PC explaining 13.3% of variance, suggesting a relatively low-dimensional representation through the population in response to the continuum of objects and scenes. To visualize this geometry, we employed both a 2D multidimensional scaling (MDS) and in 3D principal component (PC) space (Figure 4A). In both cases, a smooth pattern along spatial scale was clearly shown; in this population code, these object-to-scene conditions trace out a surprisingly linear trajectory along the major axis of variation. These analyses indicate that the depicted spatial scale of a view is represented systematically and parametrically in this larger population level code.

To further explore the relationship between voxel tuning and the resulting structure and dimensionality of the population-level representational geometry, we performed simulations with idealized voxel tuning profiles. First, we modeled pure ramp-shaped tuning, with two pools of simulated voxels tuned to each extreme end of the object-to-scene continuum (Figure 4B, see methods). Second, we modeled narrow Gaussian tuning along the continuum, with different units tuned to different scales along the continuum (Figure 4C).

The resulting representational geometry of these simulated voxel populations revealed clear differences between the two tuning motifs. For the ramp-shaped tuning, we observed an RDM with a broad diagonal and the first PC explaining 100% of the variance, with conditions mapping at a perfectly linear trajectory in the population code. In contrast, the narrow Gaussian tuning showed an RDM with a sharper and clear diagonal, a higher dimensional code (i.e., more curved eigenspectrum), with a curved (cardioid-like) trajectory in the population code. Qualitatively, the ramp-shaped tuning shows more similar signatures to the brain data than the narrow Gaussian tuning encoding scheme.

However, when we increased the width of Gaussian (i.e., broader tuning), the representation geometry becomes increasingly similar to the ramp-shaped tuning simulation. This is because wide Gaussians with peak tuning closer to the extreme ends of the continuum start to resemble the opponent ramp tuning functions, as shown in the RDM visualization (Figure 4D). Thus, in practice it is difficult to distinguish these two coding schemes, even though they have very different theoretical implications (Suzuki et al., 2005; Calder et al., 2008). However, one apparent difference between them is that with the wide-Gaussian model, the eigenspectrum does not collapse to 1 dimension, as there are at least some voxels with a preference at the middle of the continuum which cannot be modeled by combinations of two ramp shaped functions. This qualitative pattern in the eigenspectrum is actually slightly more similar to the vOTC voxels, and is perhaps consistent with the fact that while most of the voxels showed ramp-shaped tuning with peaks at the extreme, we also found a small cluster of voxels with peak tuning at an intermediate scale.

Thus, our exploratory simulations overall reveal that monotonically increasing/decreasing tuning functions (either from the ramp-shaped or wide Gaussian) are important for a parametric representation of depicted spatial scale in the population code. By having these two pools of voxels that are tuned to the opposite extremes, the object-to-scene continuum is clearly represented as linear trajectory through this large-scale population of voxels, parametrically as a function of depicted spatial scale. These simulations also provide evidence against a coding scheme in which different neural sub-populations are tightly tuned with different peaks along this depicted scale dimension; however, they also leave the open possibility that there may be wider tuning curves, or some other encoding scheme, as none of the simulations qualitatively capture the smooth curve of the eigenspectrum and the blockier similarity structure evident in the brain data RDM.

## DISCUSSION

In this study, we investigated how the depicted spatial scale of a visual scene is represented in the human visual cortex. Using rich virtual environments with controlled spatial layouts, we tested a range of depicted spatial scales from a close-up object view to a full scene view, densely sampling views in between. Considering voxel tuning, we found evidence for large swaths of cortex with opposing ramp-shaped tunings, with a smaller region with weak intermediate scale tuning. Considering the corresponding implications for the structure of multi-voxel patterns across this entire cortex, we observed that object-to-scene continuum was represented smoothly in the population code. Our simulations confirmed that in order to induce such representational geometry, it is crucial for individual voxels to show monotonic increasing or decreasing voxel tunings, which are the tuning profile we observed in the response profile clustering. Altogether, our results show that the depicted spatial scale of a view is represented parametrically in a large population of voxels, rather than with a smooth continuum of response preferences along the cortical surface.

### Independent vs Competitive Populations

Here we posit two populations with opposing ramp-shaped responses across the object-to-scene continuum. What features drive responses in each of these populations, and how do they relate to each other? One possible account is that each ramp-shaped population is coding complementary aspects of a visual environment in parallel, and *independently* (e.g., Oliva and Torralba, 2001). This account is compatible with previous works (Park et al., 2011; Harel et al., 2013), which argued that the PPA represents the spatial boundaries of a scene (e.g., walls and floors) and the LOC represents the content of the scene (e.g., textures, materials, or objects). In our data, the ramp-shaped tunings were found in two distinctive clusters in the vOTC beyond the PPA and LOC, which suggests that those responses reflect not only category-specific features that functional ROIs are sensitive to, but more general visual features. In fact, the data pattern is consistent with previous findings showing sharp dissociations along the mid-fusiform sulcus (MFS; Grill-Spector and Weiner, 2014), where there are a clear lateral to medial transitions in anatomical features such as cytoarchitecture (Weiner et al., 2014) or white-matter connectivity (Saygin et al., 2012), as well as functional responses, such as eccentricity bias (Hasson et al., 2002), animacy (Haxby et al., 2011), or real-world object size (Konkle and Oliva, 2012). Based on these, it was argued that visual information is sorted and mapped onto different sides of the MFS, as a solution to the problem of organizing high-dimensional information along the 2D cortical sheet (Grill-Spector and Weiner, 2014). Under this account, the ramp-shaped responses across depicted spatial scale in these regions are a consequence of the way different visual features (or combinations of them) naturally co-vary with the depicted spatial scale, rather than the direct result of sensitivity to scale information per se.

An alternate possibility is that the responses of these two populations are directly competitive and inter-related. For example, there might be connectivity that ensures that when one population is activated high another population is suppressed. In fact, there is some evidence for such competitive relationship (Mullin and Steeves, 2013). When the LO was disrupted using transcranial magnetic stimulation (TMS), the response in the PPA increased for scene stimuli, suggesting inhibitory connections between them. It is plausible that such connection is present across a larger populations, not just between the PPA and LO. Although we cannot tease apart these two accounts here, it may be possible to test them by independently manipulating different aspects of images. For example, we could create identical environments but without much object content while leaving the spatial structure intact. The first account predicts that responses in only one of populations will be affected (e.g., object-processing systems), whereas the second account predicts that both populations will be affected due to their connections. However, further research is needed to clearly separate and characterize different aspects of visual features in scenes.

### Implications for read-out of depicted scale information

Our results showed that coupling two opposing ramp-shaped tuning functions allows for a fine-grained representation of scale at the population level. In fact, other studies have proposed a few computational advantages of this kind of opponent coding (Regan and Hamstra, 1992). First, it allows for simple and efficient read-out by providing linearly separable representations (Guigon, 2003; Grill-Spector and Weiner, 2014). Second, the read-out is robust to baseline activation changes (e.g., from adaptation), compared to having a single monotonic tuning function (Lee et al., 2020), as an exact position can be computed as a balance of two populations. Third, fine-grained level discrimination is possible; partially overlapping tuning functions allow interpolations between discrete feature values (e.g., peaks of tuning).

Further, the opposing ramp-shaped tunings have been highlighted as a principle encoding scheme underlying geometric shape coding or facial feature coding (Freiwald et al., 2009; Kayaert et al., 2005). For example, in the face patch of macaque monkeys, cells were tuned to different subsets of facial features (e.g., face aspect ratio, inter-eye distance, iris size, etc), rather than holistic exemplars (whole faces). The tuning to individual features showed the peak response at one extreme and graded response to a minimal level at the opposite extreme. With a set of such tunings to different features, a facial identity can be constructed (e.g., linear combination; Chang and Tsao, 2017). Our data are consistent with this kind of logic, but at a larger cortical scale (spanning the entire ventral occipitotemporal cortex) for the dimension of depicted spatial scale, which is a crucial feature for scenes.

### Connections with related work

To what extent our results are consistent with previous studies that asked similar questions? First, we found a small cluster of voxels showing preference for the intermediate spatial scale, in addition to the large clusters with ramp-shaped tuning. The strength of preference was relatively weak, but it was consistently observed across several analyses. This intermediate-preferring cluster might be consistent with reachspace-preferring regions (Josephs and Konkle, 2020). However, the anatomical locations of these reachspace-preferring regions were not clearly mapped onto the intermediate-preferring cluster in the current study. It is partly because many voxels in the reachspace-preferring region did not survive the voxel selection threshold in our data. Thus, we cannot draw a firm conclusion here (see Supplemental Figure 5 for details).

Second, our results provide support for the opponent coding model in response to the depicted spatial scales, rather than the multi-channel model, implied by Peer and colleagues (2019). They argued for voxel-level Gaussian tuning in representing spatial scale. Do our results contradict their findings? On the surface, two studies may seem similar, but there are some fundamental differences. First, there were no visual stimuli presented, except the written words of objects for the task. Thus, the representation of spatial scale captured in Peer et al. (2019) could be much more abstract or modality-independent compared to the representation of visually depicted spatial scale in the current study. Second, the task in our study was passive viewing of visual scenes, whereas the task in Peer et al. (2019) was judgments of distance between objects. Finally, the range of spatial scales used in Peer et al. (2019) is much larger than our study, e.g. their closest spatial scale level was indicated by the word ‘room’, followed by ‘building’, while in our study, the farthest condition was a view of an entire room. So, it is possible that a whole range of our tested spatial scales falls in the first level of conditions in Peer et al. (2019), not allowing us to see the change at larger spatial scales along the cortical anterior-posterior axis. Overall, there could be possible explanations for discrepancy between our results and Peer et al. (2019), but we are limited to make direct comparisons here due to inherent differences between two studies.

### Conclusion

In summary, we examined how the human visual cortex represents the object-to-scene continuum, in which the depicted spatial scale was continuously varied in visual scenes. We did not find any evidence for a topographic map on the cortex, which would be driven by a gradual shift in the peak of activation at individual voxels. Instead, we found a smooth representation of the object-to-scene continuum in a population of voxels, which showed two opposite ramp-shaped tunings. One important thing to note is that the object-to-scene continuum tested in our study is the pictorial description of spatial scale, which was achieved by adjusting the field of view as well as viewing distance. However, in natural viewing condition, the vergence of the eyes on an object changes the human field of view much less dramatically than implied here, and we have access to the full-field of view without an image border. Thus, one promising future direction would be testing the perception of spatial scales in full-field viewing, allowing far-peripheral vision to be engaged in processing the content of central vision.

## Methods

### Participants

Twelve participants (5 females, age: 20-38 years) with normal or corrected-to-normal vision were recruited from the Harvard University community. All participants gave informed consent and were financial compensated. All procedures were approved by the Harvard University Human Subjects Institutional Review Board.

### Stimuli

Computer generated (CGI) environments were generated using the Unity video game engine (Unity Technologies, Version 2017.3.0). We constructed twenty indoor environments, reflecting a variety of semantic categories (e.g., kitchens, bedrooms, laboratories, cafeterias, etc.). All rooms had the same physical dimensions (4 width x 3 height x 6 depth arbitrary units. in Unity), with an extended horizontal surface along the back wall, containing a centrally-positioned object. Each environment was additionally populated with the kinds of objects typically encountered in those locations, creating naturalistic CGI environments.

Images spanning a continuum of distances from the central object were captured from each environment (1024 × 768 pixels, approximately 20 visual angles wide), ranging from a close-up view of the object to a far-scale view, which included the whole room. Images were generated by systematically varying the location of the camera (hereafter “Position”) along 15 log-scaled points arrayed from the “front” to the “back” of the room (i.e., from right in front of the central object to across the room from it, Figure 1A). Close-up views were captured with a smaller camera field of view (FOV), so that only the central object appeared in the frame, and the FOV increased logarithmically with each step away from the object. The camera angle was parallel to the floor plane for far-scale views, and was gradually adjusted downward for closer positions, so that the central object was always at the center of the image Figure 1B). These camera parameters were used for all 20 environments, yielding 300 unique stimuli (20 environments x 15 positions). See https://osf.io/hcmgk/ for all stimuli.

### Experimental Design

The main experiment consisted of 10 runs. Each run was 6.2 min in duration (186 TRs), and used a standard blocked design, with 15 conditions presented twice each run. Each condition block was 6s, in which five images from one condition were presented (e.g., 5 different environments viewed the same position), and was always followed by a 6s fixation period. Within a condition block, each image was presented for 800 ms, followed by a 300 ms blank screen. The presentation order of blocks in each run was pseudo-randomized as follows. Fifteen conditions within an epoch were randomized twice independently and concatenated with a constraint that the same condition cannot appear in two successive blocks. The CGI environments were randomly divided into two sets (Environment Set A and B) and used for odd and even runs. These stimuli sets were kept the same within each participant. Participants performed a red frame detection in which they pressed a button whenever there was a red frame surrounding the stimulus. The red frame appeared once in a block, in a random position among 5 images. Participants were instructed to pay attention to both spatial layout and objects.

Two functional localizer runs were performed independent of the main experimental runs, each 6.9 min (208 TRs). In each run, participants saw blocks of faces, objects, scenes, and reachspaces images (Josephs and Konkle, 2020), while performing one-back repetition detection task. In each condition block (8 sec), seven unique images were selected and one of those images was randomly chosen and repeated twice in a row. Participants were instructed to press a button when they saw the repeated image. In each trial, an image was presented for 800 ms, followed by a 200 ms blank screen. Ten blocks per condition were acquired within a run. The presentation order of blocks in each run was randomized within each epoch. One epoch consisted of one block from each of four conditions and one fixation block (8 sec). This procedure was repeated ten times and the block orders were concatenated across the epochs.

Finally, two additional retinotopy runs were performed, each 5.4 min (162 TR). This protocol consisted of 4 conditions: horizontal bands (∼22º x 1.7º), vertical bands (∼1.7º x 22º), central stimulation (radius ∼ 1.2º to 2.4º), and peripheral stimulation (radius ∼ 9.3º to the edges of the screen). The horizontal and vertical bands showed checkerboards which cycled between states of black-and-white, white-and-black, and randomly colored at 6hz. The central and peripheral rings showed photograph fragments which cycled between patterns of object ensembles (e.g., buttons, beads) and scene fragments (Cant and Xu, 2012; Zeidman et al., 2018). Each run consisted of 5 blocks per condition (12 sec block), with five 12 sec fixation blocks interleaved throughout the experiment. An additional 12 sec fixation block was added at the beginning and the end of the run. Participants were asked to maintain fixation and press a button when the fixation dot turned red, which happened at a random time once per block.

The stimuli presentation and the experiment program were produced and controlled by MATLAB and Psychophysics Toolbox (Brainard, 1997; Pelli and Vision, 1997).

### fMRI Data Acquisition

All neuroimaging data were collected at the Harvard Center for Brain Sciences using a 32-channel phased-array head coil with a 3T Siemens Prisma fMRI Scanner. High-resolution T1-weighted anatomical scans were acquired using a 3D MPRAGE protocol (176 sagittal slices; FOV = 256 mm; 1×1×1 mm voxel resolution; gap thickness = 0 mm; TR = 2530 ms; TE = 1.69 ms; flip angle = 7°). Blood oxygenation level-dependent (BOLD) contrast functional scans were obtained using a gradient echo-planar T2* sequence (84 oblique axial slices acquired at a 25° angle off of the anterior commissure-posterior commissure line; FOV = 204 mm; 1.5 × 1.5 × 1.5 mm voxel resolution; gap thickness = 0 mm; TR = 2000 ms; TE = 30 ms, flip angle = 80°, multi-band acceleration factor = 3).

### fMRI Data Analysis and Preprocessing

The fMRI data were analyzed with BrainVoyager 21.2.0 software (Brain Innovation) with custom Matlab scripting. Preprocessing included slice-time correction, linear trend removal, 3D motion correction, temporal high-pass filtering, and spatial smoothing (4mm FWHM kernel). The data were first aligned to the AC-PC axis, then transformed into the standardized Talairach space (TAL). Three-dimensional models of each participant’s cortical surface were generated from the high-resolution T1-weighted anatomical scan using the default segmentation procedures in FreeSurfer. For visualizing activations on inflated brains, the segmented surfaces were imported back into BrainVoyager and inflated using the BrainVoyager surface module. Gray matter masks were defined in the volume based on the Freesurfer cortex segmentation.

A random-effect group general linear model (GLM) was fit using BrainVoyager. The design matrix included regressors for each condition block (Position 1-15) and 6 motion parameters as nuisance regressors. The condition regressors were constructed based on boxcar functions for each condition, convolved with a canonical hemodynamic response function (HRF), and were used to fit voxel-wise time course data with z-normalization and correction for serial correlations. The extracted beta weights from this group GLM were averaged across participants for each voxel, and then taken as the primary measure of interest for all subsequent analyses. The group data was displayed on a selected participant’s cortical surface.

### Regions of Interest (ROIs)

We identified five ROIs separately in each hemisphere in each participant, using condition contrasts implemented subject-specific general linear models. Three scene-selective areas were defined using Scenes - Objects (*p <* .0001) contrast. Specifically, the parahippocampal place area (PPA) was defined by locating the cluster between posterior parahippocampal gyrus and lingual gyrus, the retrosplenial cortex (RSC) was defined by locating the cluster near the posterior cingulate cortex, and the occipital place area (OPA) was defined by locating the cluster near transverse occipital sulcus. The lateral occipital complex (LOC) was defined using Objects - Scenes (*p <* .0001) contrast. The fusiform face area (FFA) was defined using Faces - Objects (*p <* .0001) contrast. Finally, the early visual areas (EVA; V1-V3) were defined manually on inflated brain, based on the contrast of Horizontal – Vertical meridians from the retinotopy runs.

### Voxel Selection

The preference mapping and response profile clustering analyses were performed on voxels selected using the anatomical mask and the reliability-based voxel selection method (Tarhan and Konkle, 2020a). First, we manually drew a mask in BrainVoyager (21.2.0) encompassing occipito-temporal cortex, occipito-parietal cortex, and the corresponding medial part of the brain. Within this mask, we calculated split-half reliability for each voxel by correlating the betas from odd and even runs of the main experimental runs. Based on the resulting correlation map, we chose *r*=0.3 as the cutoff and selected voxels with higher reliability (Supplemental Figure 1). This voxel selection procedure was performed on a group-level in the Talairach space.

### Preference Mapping

To examine whether there is a topographic mapping of the object-to-scene continuum, we calculated a group-level preference map. First, responses to each of 15 Positions were extracted in each voxel from single-subject GLMs, then averaged over subjects. For each voxel, a condition showing the highest group-average response was identified as the preferred condition. The degree of preference was computed by taking the response differences between the most preferred condition and the next-most-preferred condition. To visualize these response topographies, we colored each voxel with a color hue corresponding to the preferred condition, with a color intensity reflecting the degree of preference. This preference map was projected onto the cortical surface of a sample participant. Similar preference mapping procedures have been used in the studies examining a large-scale cortex organization (Orlov et al., 2010; Konkle and Caramazza, 2013; Long et al., 2018; Josephs and Konkle, 2020; Masson and Isik, 2021).

### Response Profile Clustering

We used a data-driven neural clustering method to see whether there are clusters of voxels responding similarly over the conditions. Using MATLAB’s implementation of the *k*-means algorithm with the correlation distance, voxels were grouped based on their response profile similarity (10 replicates, 500 max iterations). To measure the similarity, the response profiles for all voxels were first transformed to have zero-mean and unit length, and then correlation was used as the distance metric. We chose to use the correlation in order to compare relative response magnitudes across the conditions, rather than differences in overall response magnitudes, which may be sensitive to a voxel’s anatomical location and signal measurement related reasons (e.g., proximity to head coils). The output of this analysis was a clustering assignment for each voxel, with a cluster centroid response profile, which reflects the average normalized profile for all voxels included in the cluster. To visualize the clustering solution, we created cortical maps in which all voxels assigned to the same cluster were colored the same. For visualization purposes, we chose these colors in a way that clusters with more similar response profiles were more similar in hue. To do so, we submitted the cluster response profiles to a multidimensional scaling algorithm using a correlation distance measure, placing similar cluster centroids nearby in a three-dimensional space. The 3D coordinates of each point were re-scaled to be within [0 1], and then used as the Red-Green-Blue color channels for the cluster color.

To determine the number of cluster solutions, we computed a range of *k*-means solutions varying the number of clusters (*k*) from 2 to 20. We chose the final *k* based on the cluster center similarity (Supplemental Figure 3A), which measures how similar the cluster centers are to one another on average. We considered solutions with low cluster center similarity, because if two clusters are very similar to each other, it is difficult to draw any meaningful differences or interpret them.

For assessing the stability of clustering solution, we conducted two additional analyses. First, we assessed the reliability of the clustering solution across different subsets of stimuli (Supplemental Figure 3C). The data were split into two based on the stimuli (Environment Set A vs. Environment Set B), and the identical clustering algorithm was ran separately for each data set. Then, we quantified how well those two clustering solutions converged, using a signal detection method (Konkle and Caramazza, 2017). The d-prime (d’) score was computed by comparing the voxel cluster assignments between two solutions. For each solution, we created a matrix (voxels x voxels), in which the value is 1 if the voxels were assigned to the same cluster and 0 if the voxels were assigned to different clusters. Then, Hit rate was calculated as the percentage of voxel pairs that were assigned to the same cluster in one solution that were also assigned to the same cluster in another solution; and False Alarm rate was calculated as the percentage of voxel pairs that were not assigned to the same cluster in one solution but were assigned to the same cluster in another solution. The sensitivity measure d’ was calculated as z(Hit)-z(FA).

Second, we assessed the stability of the clustering solutions across participants (Supplemental Figure 3B). For each iteration, the participants were randomly split into two groups, and the sensitivity of one group to predict the other was computed using the same signal detection method described above. This d’ score was measured for each of the solutions *k*=2 to 20. Then, the entire procedure was repeated over 50 iterations, and we computed the average d’ over iterations for each clustering solution *k*.

### Population-Level Analyses

This analysis examined how the object-to-scene continuum is represented as the distributed patterns across voxels in the ventral occipitemporal cortex beyond early visual cortex. We selected a set of voxels that were located within an anatomical mask of ventral occipital temporal cortex (vOTC) and the voxel-wise reliability is higher than 0.3, while excluding the early visual cortex (V1-V3). Then, we computed the distance between each condition pairs by taking 1-Pearson correlation value, which resulted in a 15 × 15 representational dissimilarity matrix (RDM). The pairwise distance relationship between conditions was also visualized using a 2-dimensional classic multidimensional scaling (MDS) function implemented in MATLAB R2020B.

To further explore the geometry of the neural code, we performed a dimension reduction technique, principal component analysis (PCA), over the selected voxels. The PCA function was implemented in MATLAB R2020B. Then, we made an eigenspectrum plot which shows the proportion of variance explained by individual principal components. Finally, we visualized how each condition is represented in 3-dimensional PC space.

### Hypothetical Voxel Tuning Simulation

To better understand the relationship between individual voxel tunings and their population-level representation, we ran simulations comparing two coding models with idealized voxel tunings: 1) multi-channel coding with Gaussian tuning, and 2) two-opponent coding with ramp-shaped tuning. For the multi-channel model, we generated simulated activations of 150 voxels based on a Gaussian distribution.

The *μ* of Gaussian distribution was randomly chosen between 1-15 (i.e., corresponding to each stimulus condition). To examine how the representational geometry changes depending on the width of Gaussian tuning, we varied *σ* from 1 to 10. For the two-opponent model, we generated simulated activations of 150 voxels that have a linear shape; half of them were decreasing (i.e., object-preferring), and another half was increasing across the continuum (i.e., scene-preferring) as a function of depicted spatial scale along the object-to-scene continuum. The slope of linear line was randomly varied between 0.05 - 0.5, and a half of them were assigned a negative value; the intercept of line was randomly sampled between 0.01 - 0.2. After creating these hypothetical activations, we extracted distributed patterns of simulated voxels for each condition and computed the pairwise distance (1-Pearson correlation). The PCA and visualization using the RDM and MDS plots were also performed as we did with the actual voxels.

## Acknowledgments

We would like to thank George Alvarez for helpful comments on the manuscript. Research reported in this publication was supported by the National Eye Institute of the National Institutes of Health under Award Number R21EY031867. The content is solely the responsibility of the authors and does not necessarily represent the official views of the National Institutes of Health. This research was carried out at the Harvard Center for Brain Science and involved the use of instrumentation supported by the NIH Shared Instrumentation Grant Program (S10OD020039). We acknowledge the University of Minnesota Center for Magnetic Resonance Research for use of the multiband-EPI pulse sequences.

## Supplementary Information

**Supplementary Figure 1:**
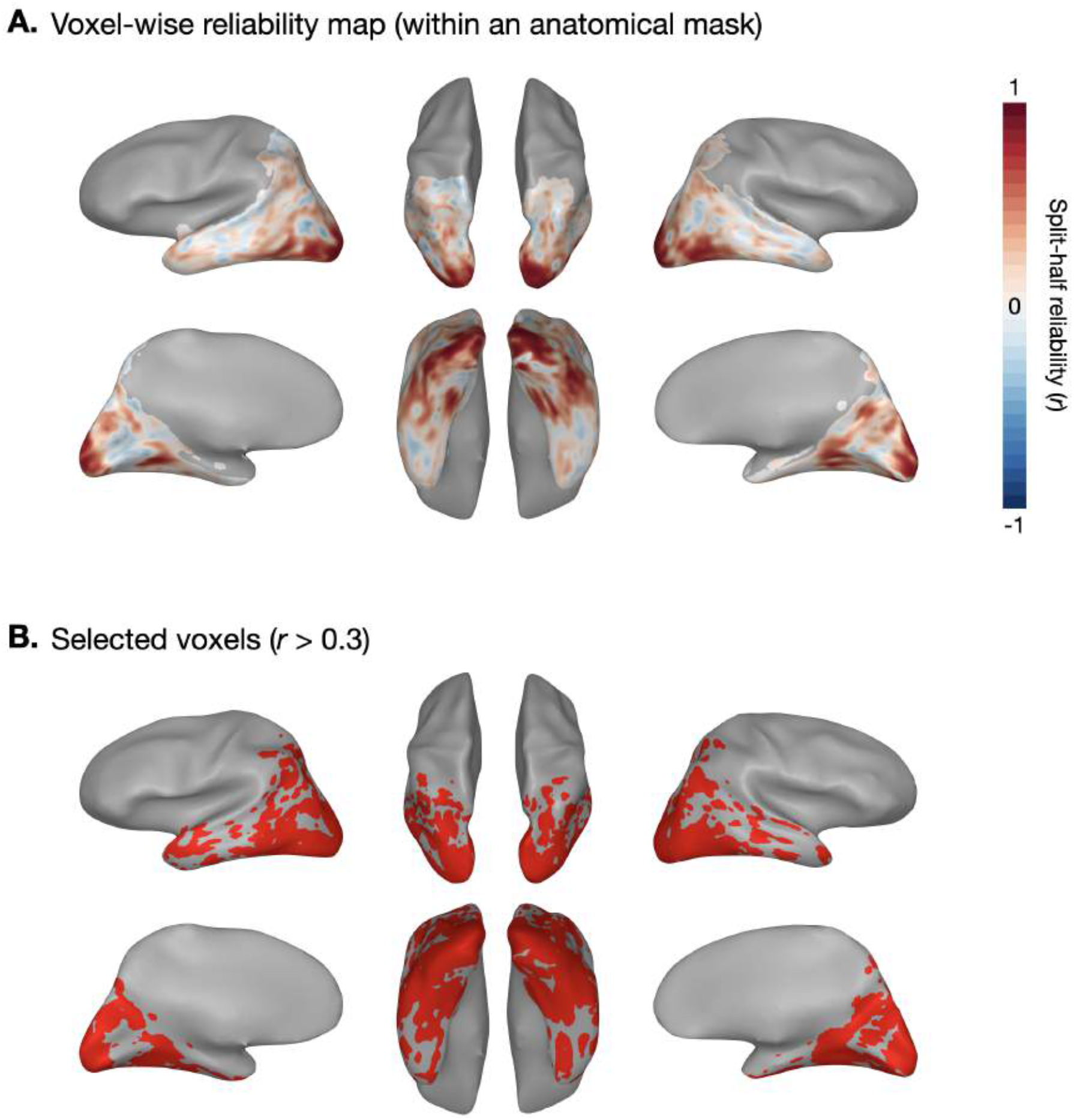
Voxel Selection. (A) First, we manually defined an anatomical mask that includes occipito-temporal cortex, occipito-parietal cortex, and the corresponding medial part of the brain. Within the mask, voxel-wise reliability was measured by correlating the betas between odd and even runs, from a group-level GLM. (B) Voxels that have higher reliability than the threshold (r=0.3) were selected for subsequent analyses. These resulting voxels are shown in red.

**Supplementary Figure 2:**
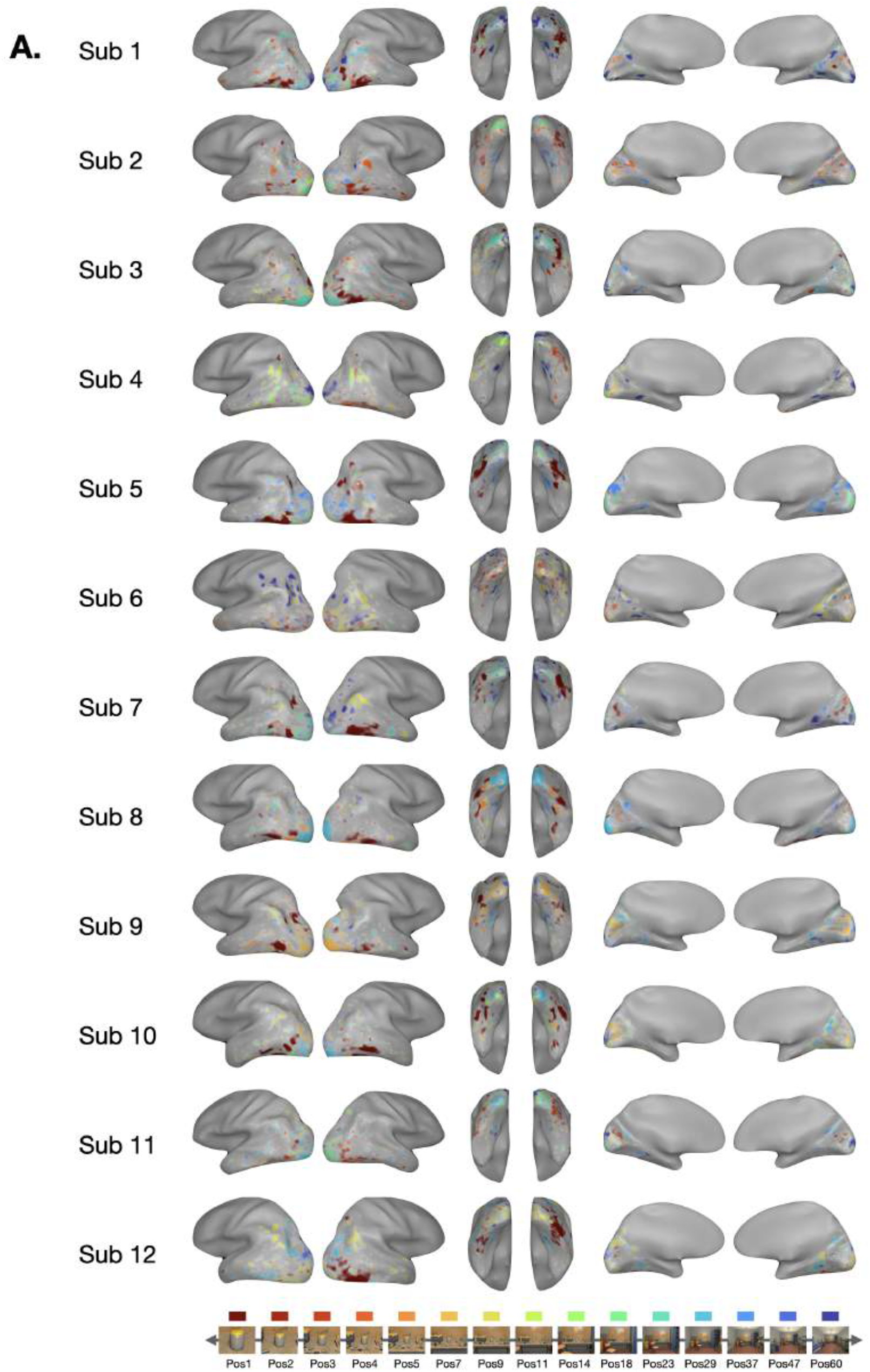
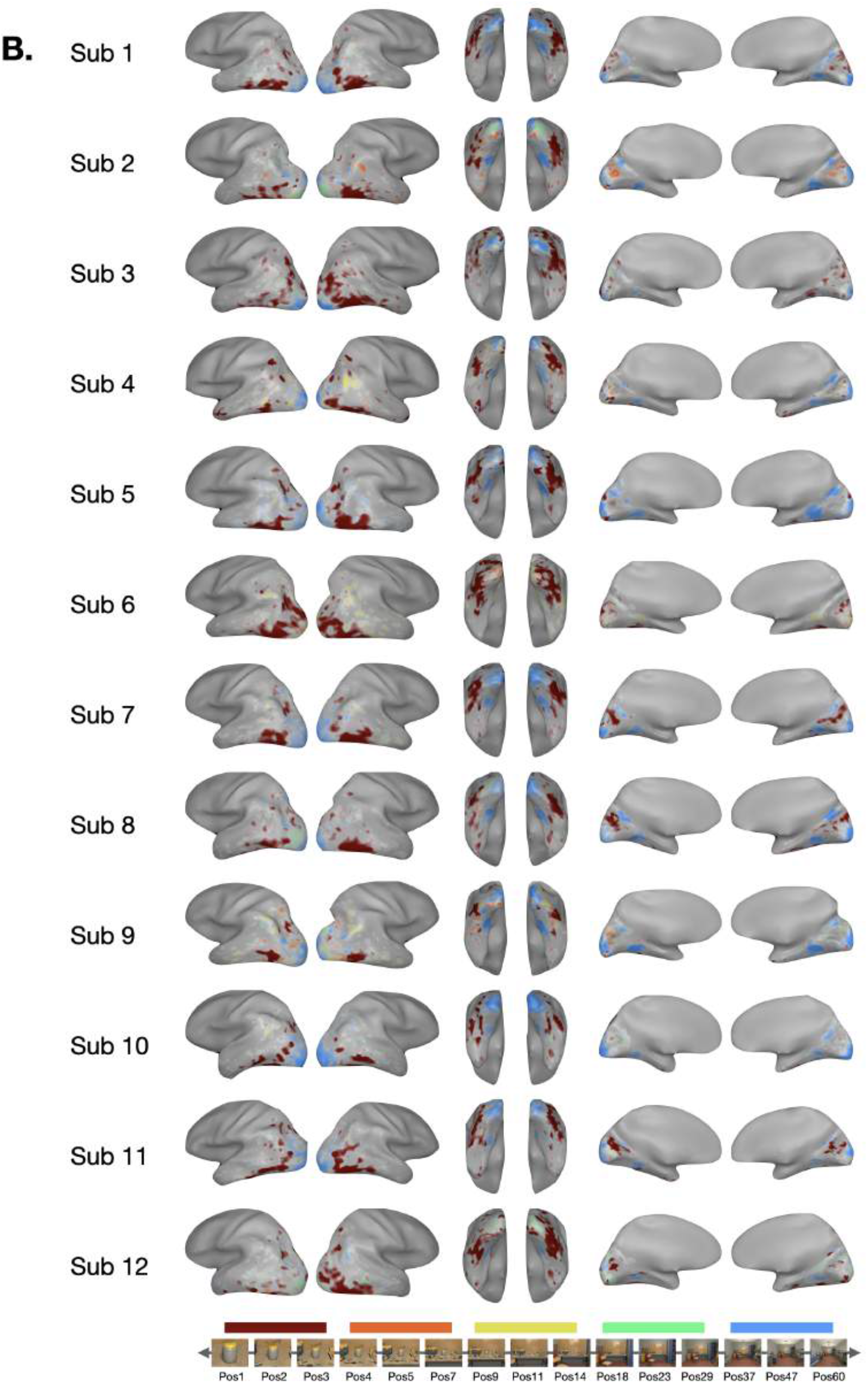
Individual Preference Maps. In addition to the group-level preference map (Figure 2), preference maps were also computed at each individual participant level. (A) Preferences for all conditions (i.e., 15 Positions) were measured, in the same manner as the group data. A lot of variation across participants was observed. (B) Voxels’ preferences were grouped and binned for 3 neighboring conditions. For example, if a voxel showed preference for condition 2, it was colored as the same color as voxels that showed preference for condition 1 or condition 3.

**Supplementary Figure 3:**
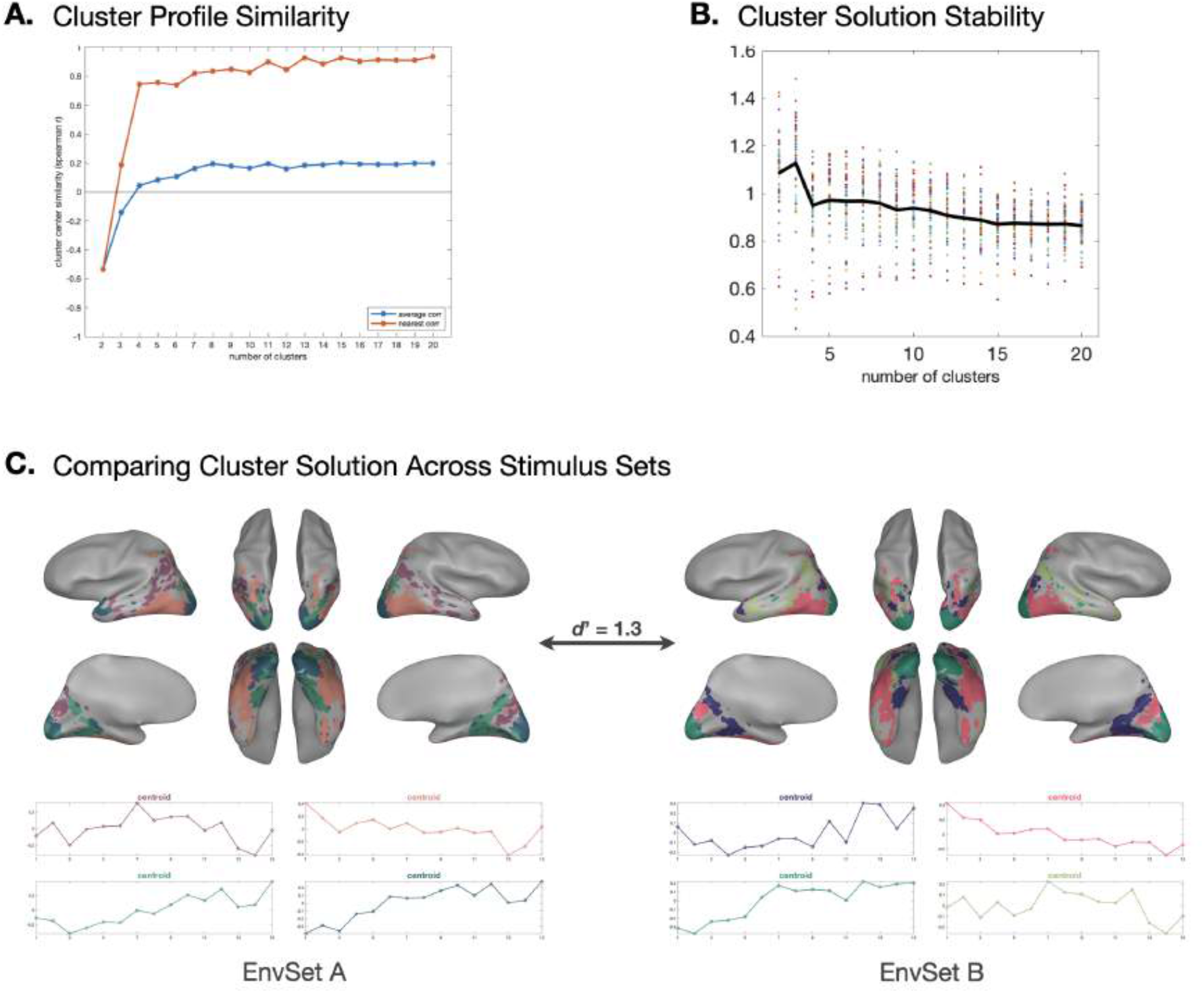
Response Profile Clustering Stability. (A) For each number of clusters (k), we measured how similar the cluster centers are to one another on average. (B) The stability of the clustering solutions were assessed across participants. For each clustering solution k, the participants were randomly split into two groups, and the sensitivity of one group to predict the other (d’) was computed. This procedure was repeated over 50 iterations, and the mean d’ was calculated for each k. (C) The reliability of the clustering solution was measured across different subsets of stimuli. The data were split into two based on the stimuli (Environment Set A vs. Environment Set B), and the identical clustering algorithm was ran separately for each data set. Then, we computed d’ by comparing the voxel cluster assignments between two solutions.

**Supplementary Figure 4:**
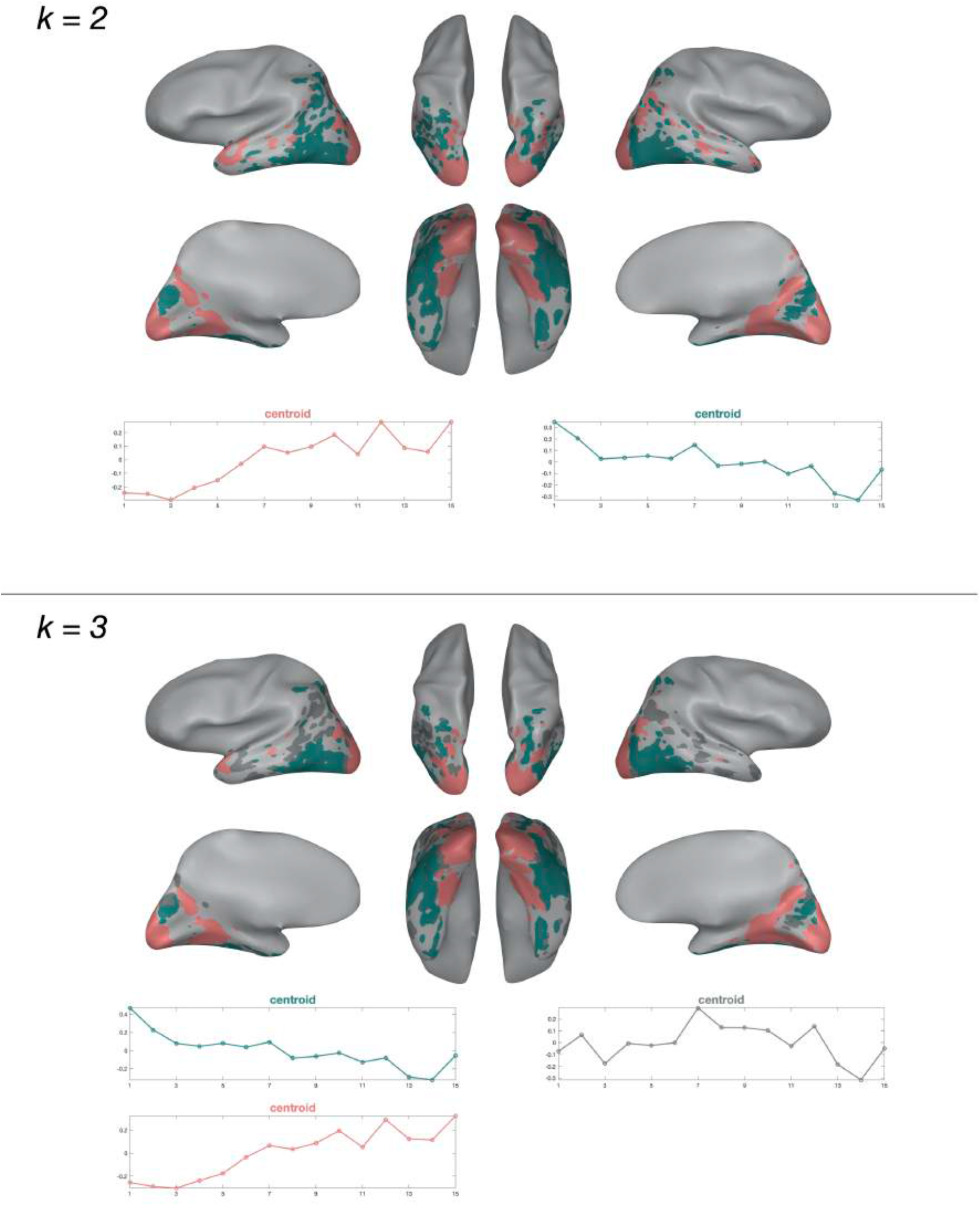
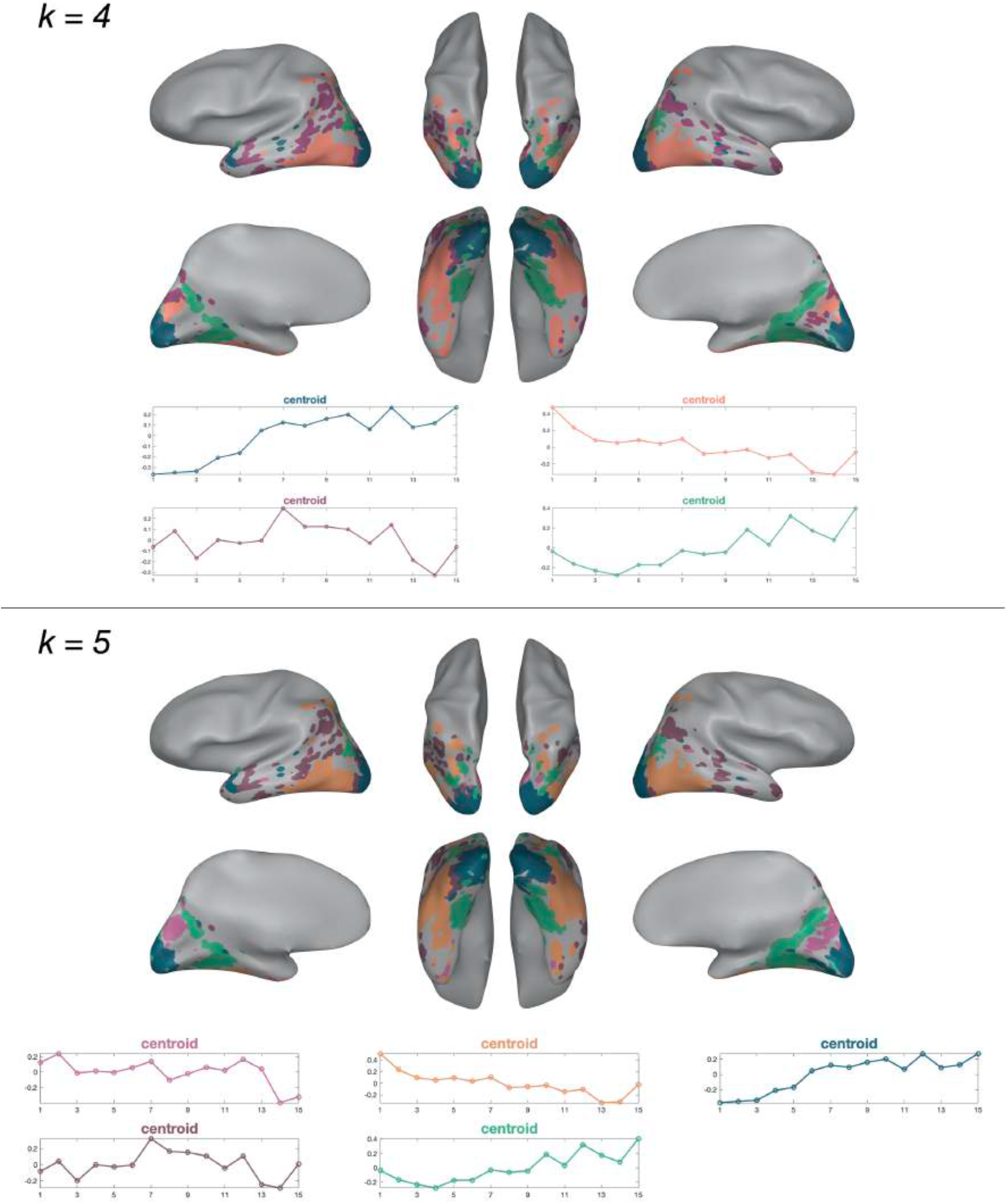
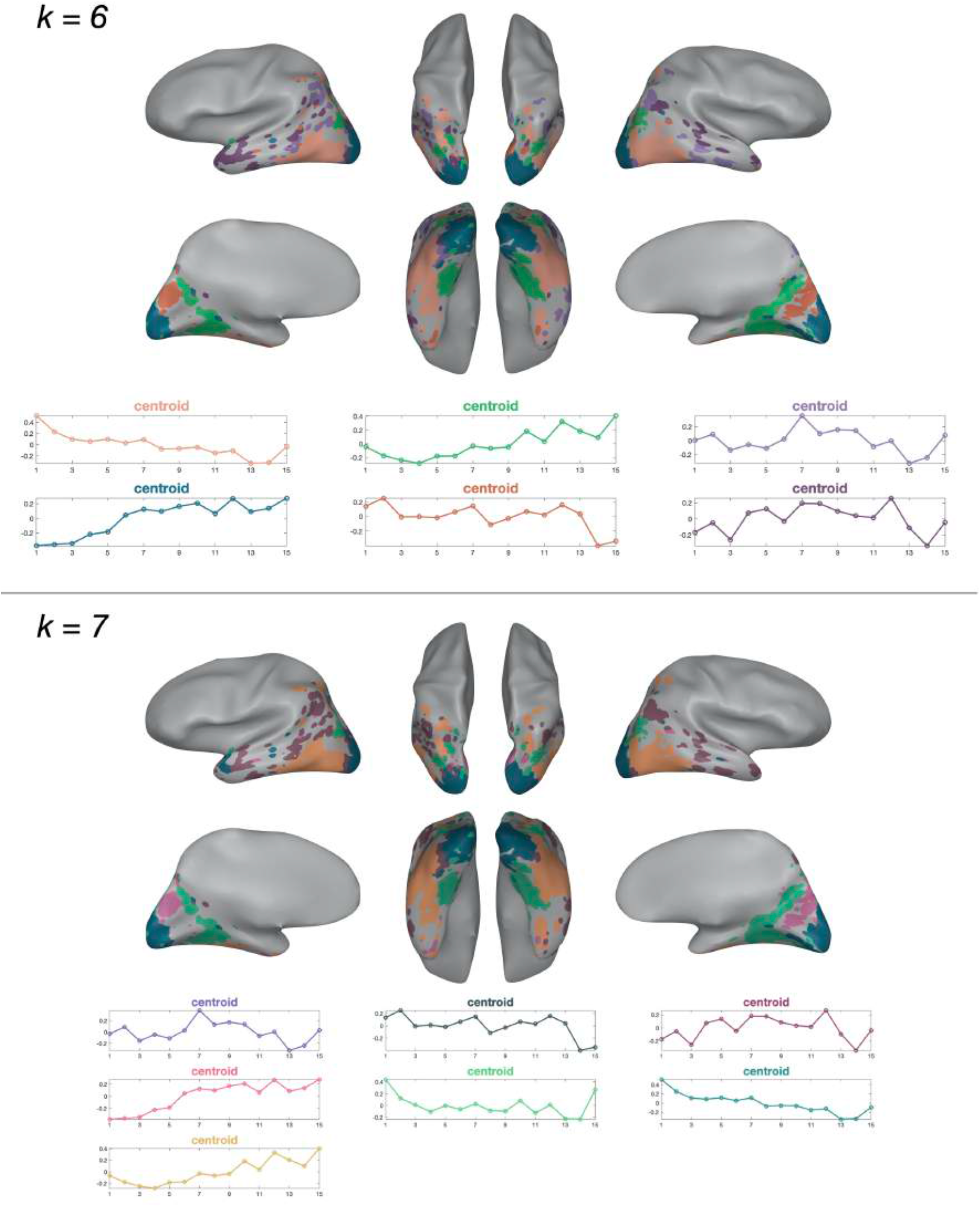
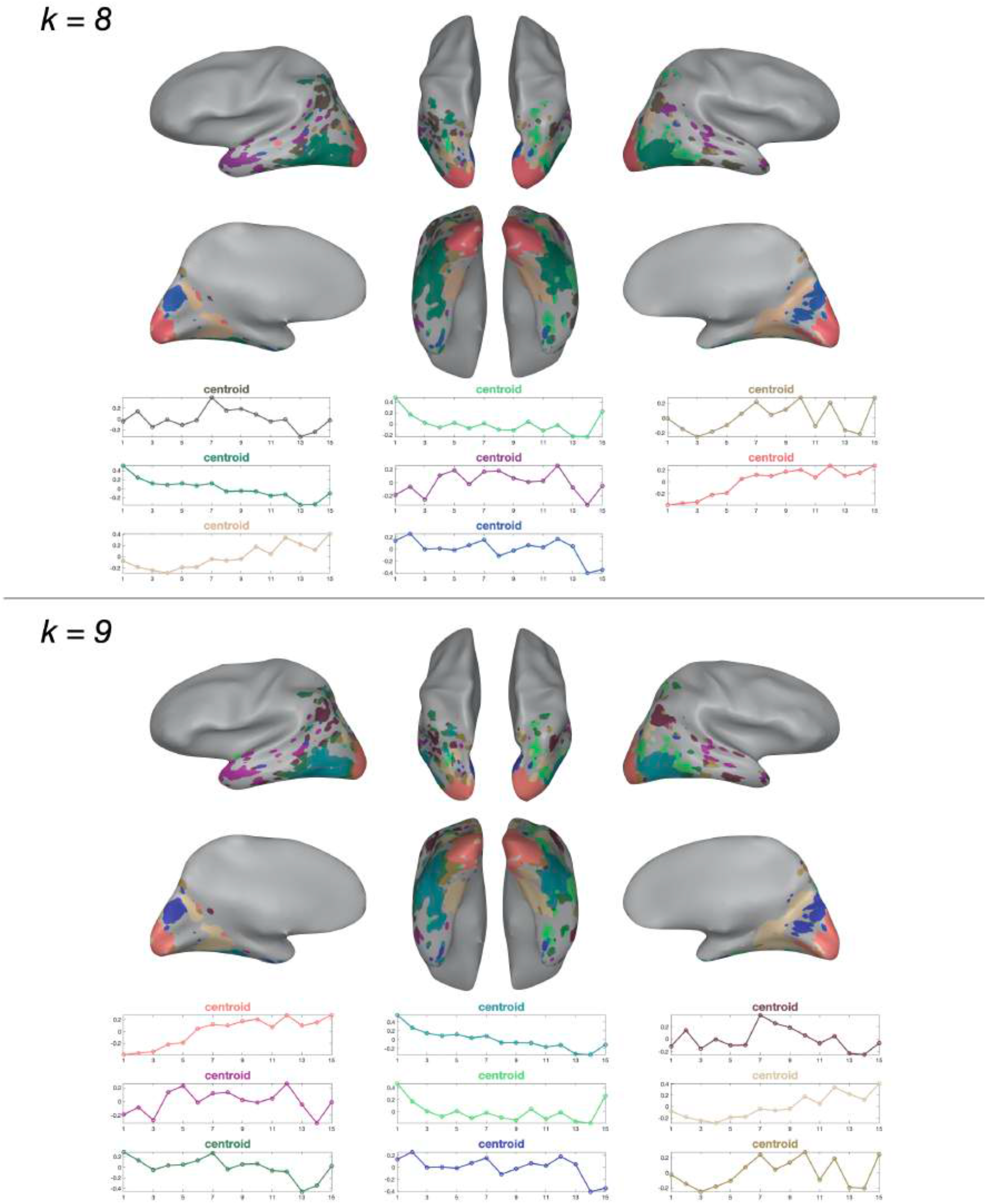
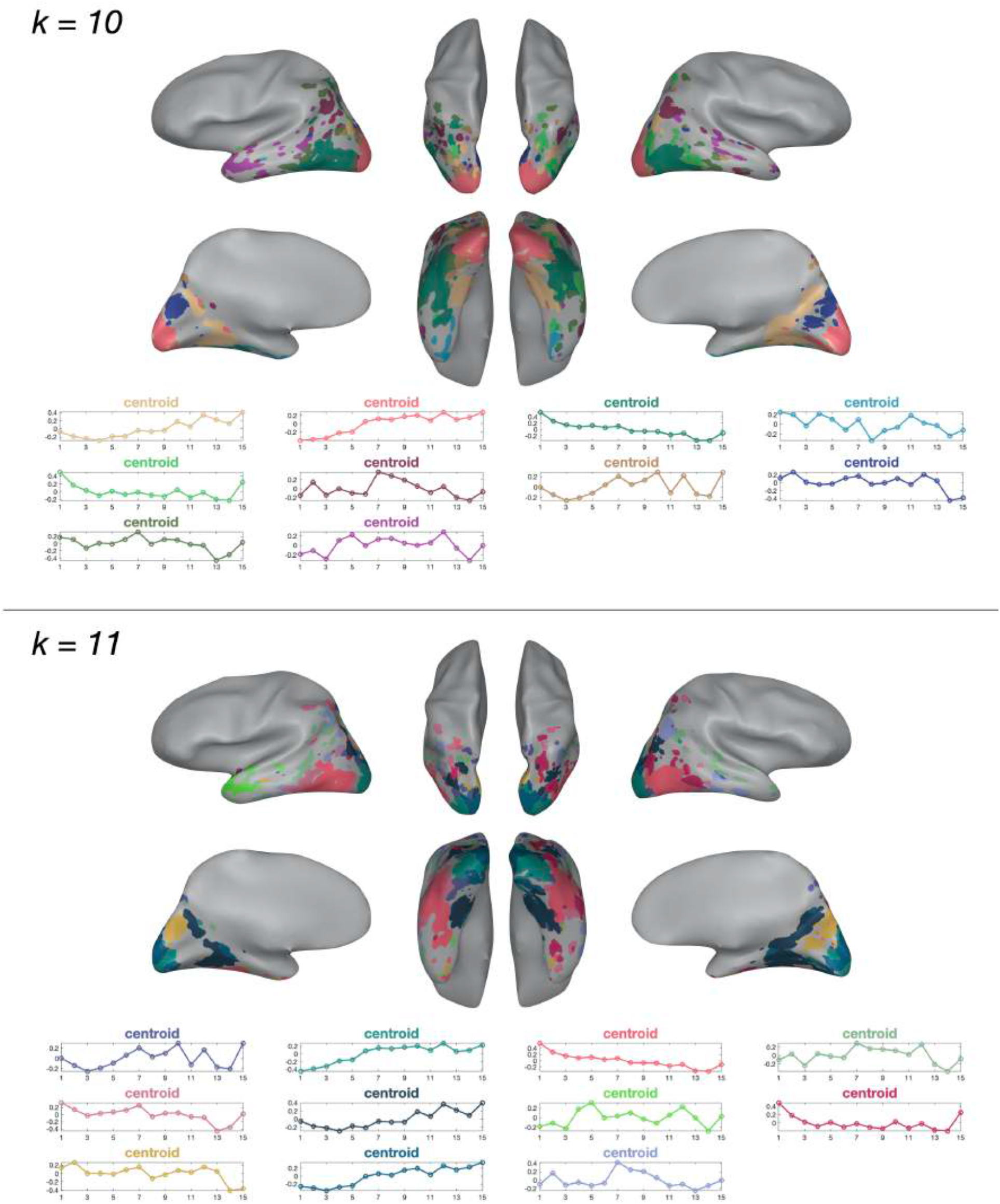
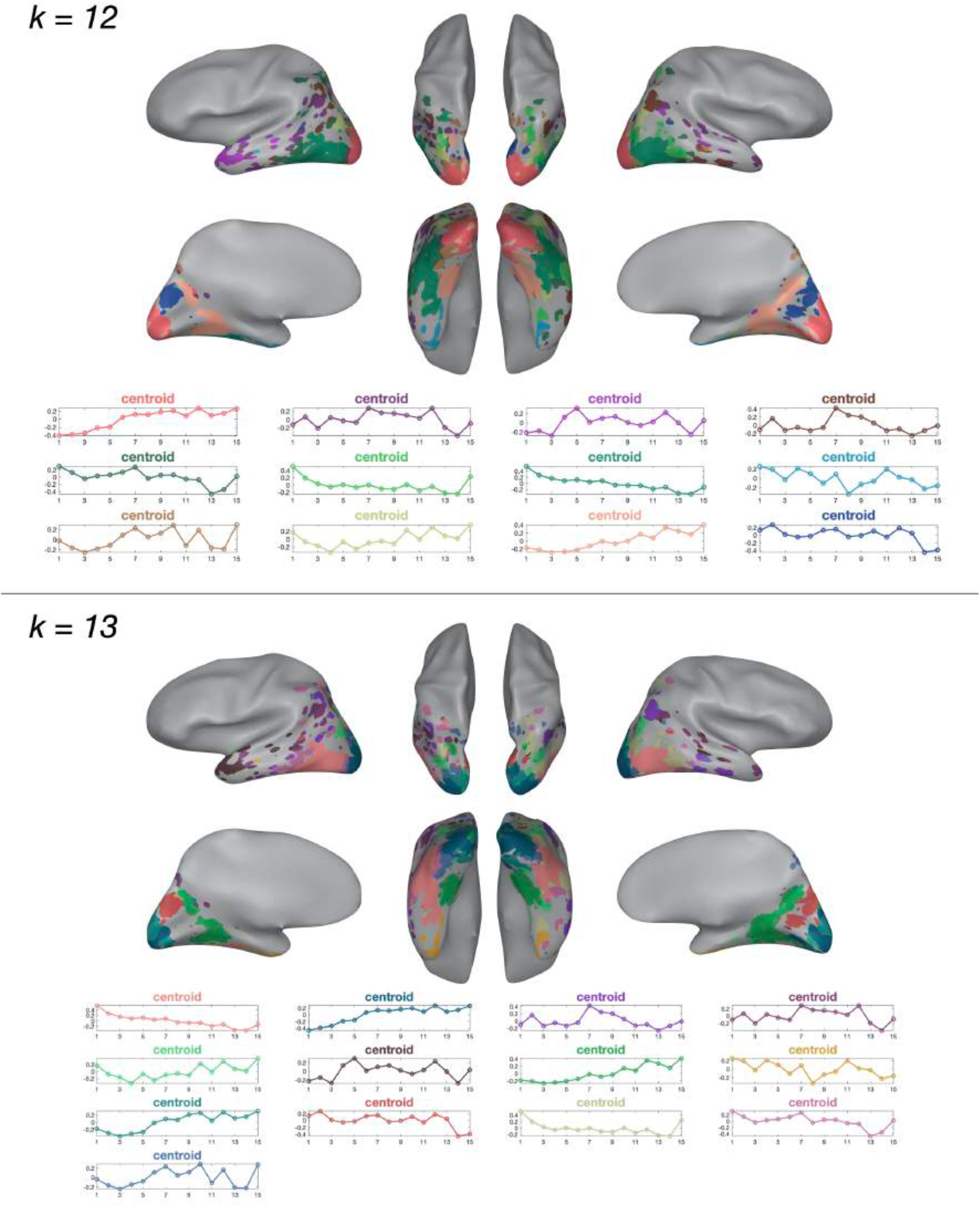
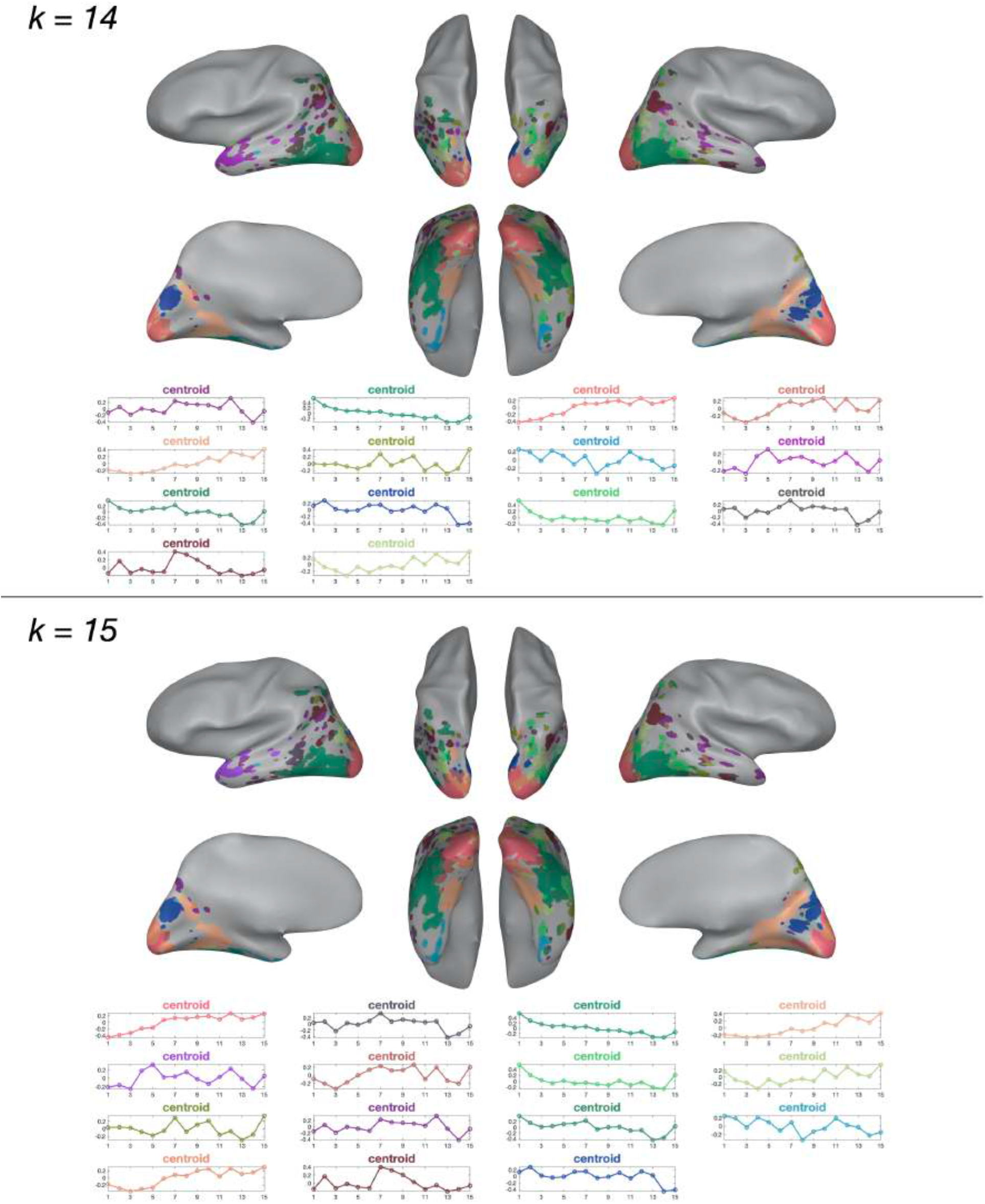
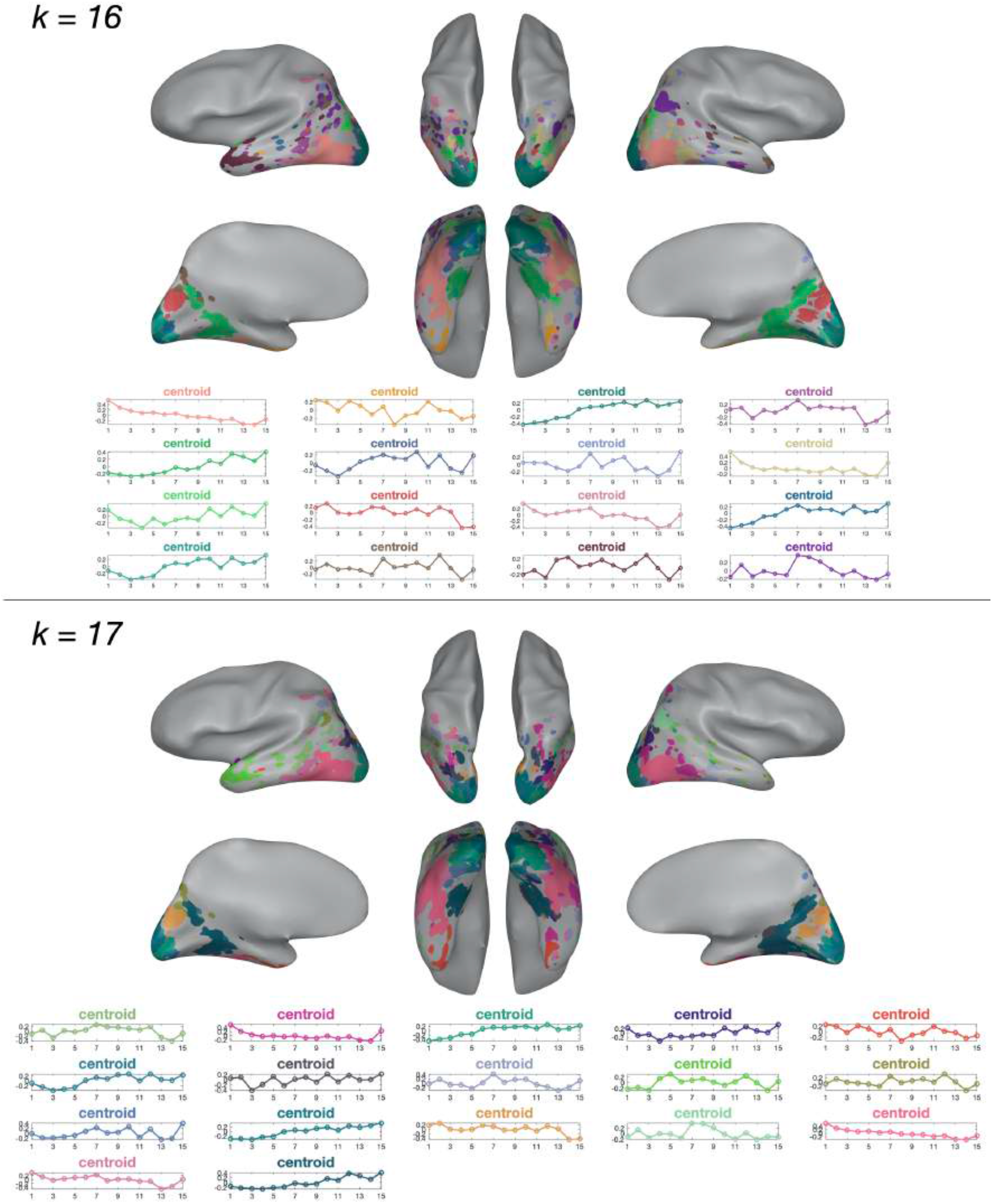
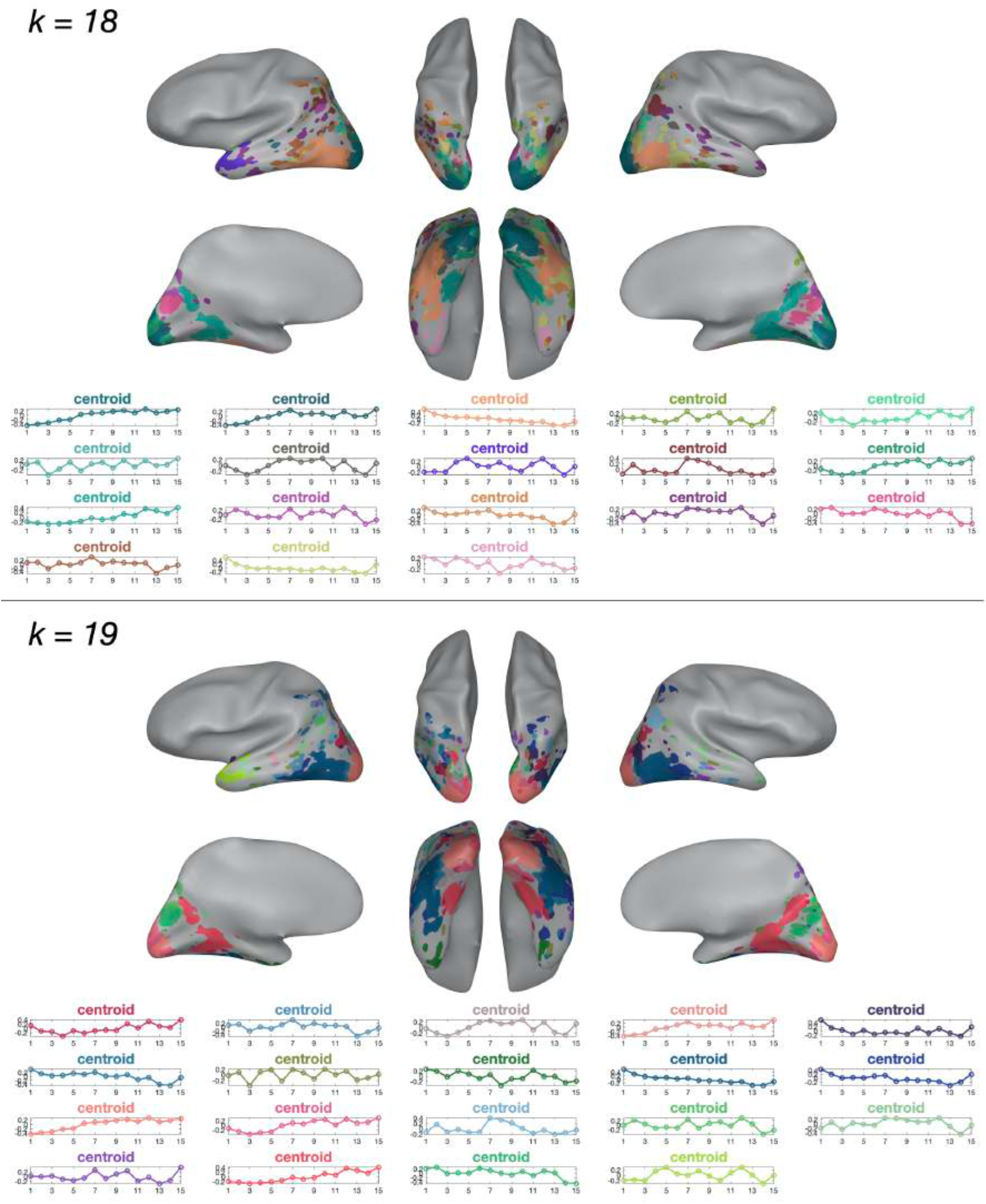
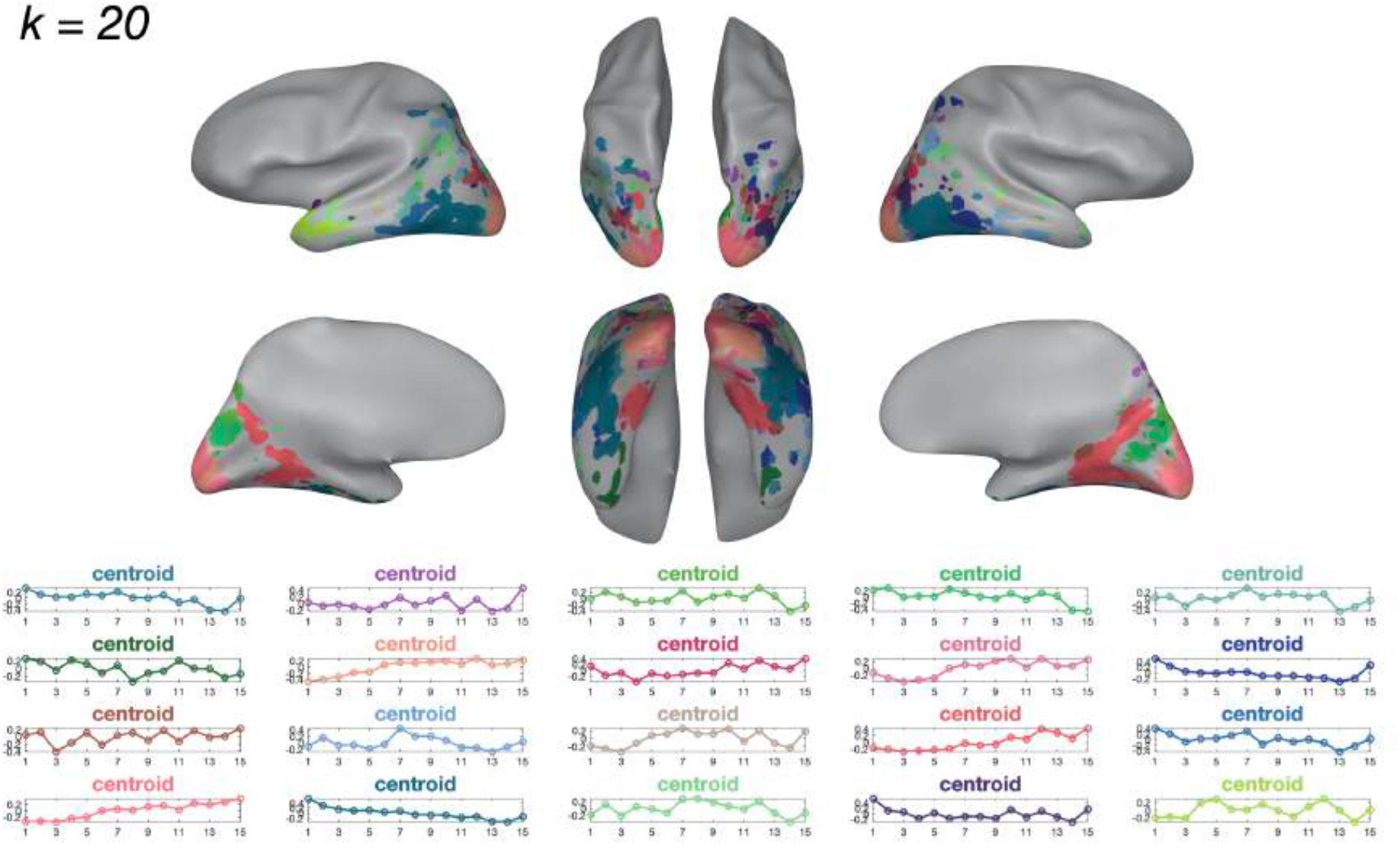
Response Profile Clustering Solutions. Clustering results for all solutions, from k=2 to k=20. Voxels assigned to the same cluster were colored the same, and we chose the colors such that clusters with more similar response profiles were more similar in hue. The corresponding cluster centroids are plotted underneath the cortical map.

**Supplementary Figure 5:**
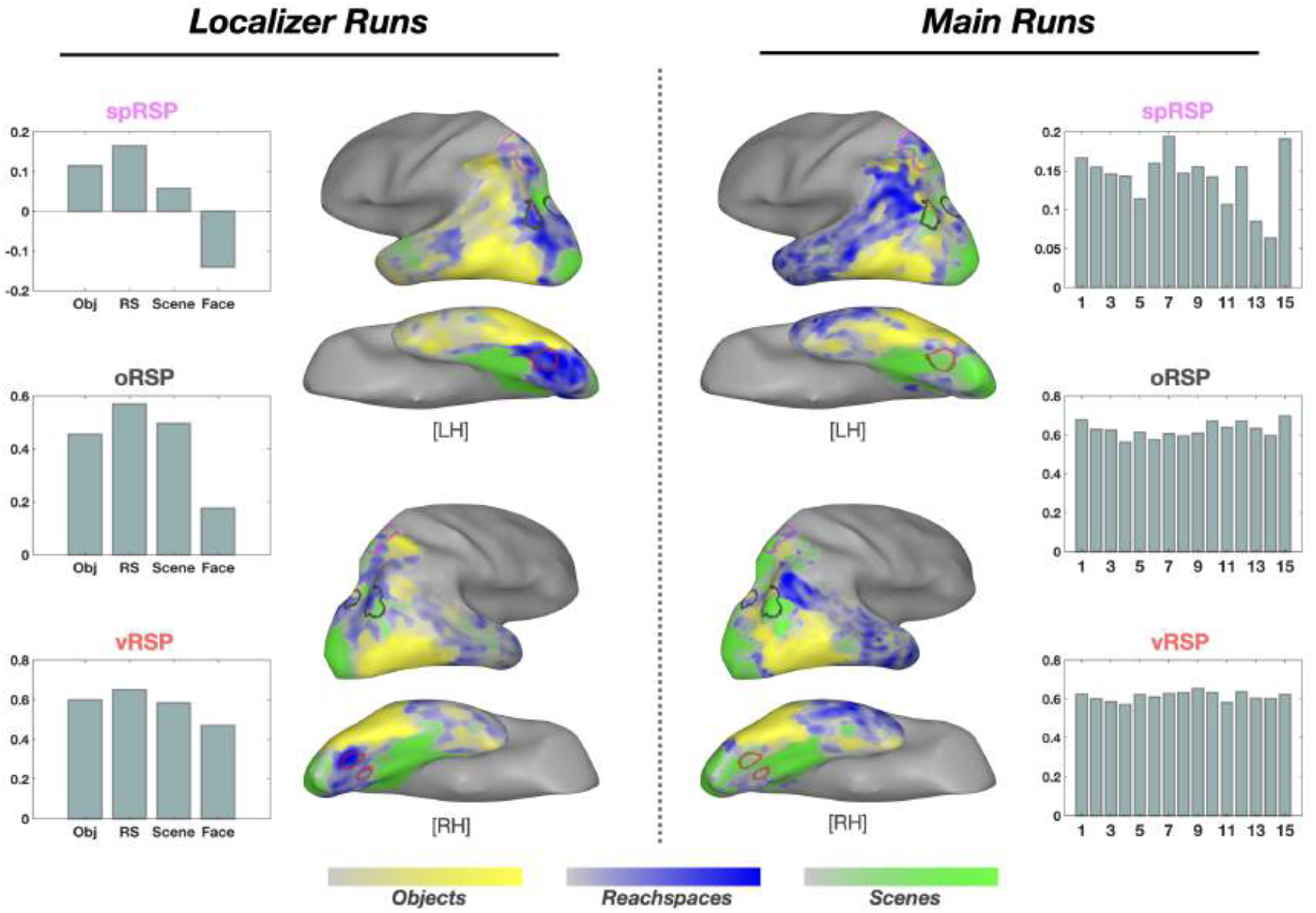
Reachspace-preferring areas. In the current study, we found some mixed evidence regarding previously reported reachspace-preferring areas (Josephs and Konkle, 2020). To directly compare our data to the previous study, we obtained three ROI masks (”protoROIs”) from the Josephs and Konkle (2020), in the superior parietal cortex (spRSP, pink), the occipiotal cortex (oRSP, black), and the ventral visual cortex (vRSP, red). First, we examined how these protoROIs responded to our stimulus conditions, separately for localizer runs and main runs. In the localizer, all 3 protoROIs showed the highest activation to the reachspaces condition compared to objects, scenes, or faces (bar graphs, left panel). In the main runs, a similar trend was shown at the superior parietal region (spRSP), but not at the occipito-parietal region (opRSP) or the ventral region (vRSP; bar graphs, right channel). Next, we computed 3-way preference maps (objects, reachspaces, and scenes) with the current study data and compared anatomical locations of the resulting reachspace-preferring cortex to the protoROIs. As the main runs did not have explicitly labeled reachspace conditions, we considered conditions 5-7 (i.e., Position 5, 7, and 9) as the reachspaces, condition 1 (Position 1) as the objects, and condition 15 (Position 60) as the scenes. Then we extracted data corresponding to those conditions and performed the preference mapping procedure. This analysis was performed on all voxels within our group-level anatomical mask (Supplementary Figure 1), without the voxel-reliability thresholding step. Considering the maps from the localizer runs, we observed close correspondence to the maps from Josephs and Konkle (2020). Additionally, most of the protoROIs showed good correspondence with the reachspace-preferring cortex. However, the maps from the main runs showed different topographies from both the localizer run and the preference maps in Josephs Konkle (2020). The discrepancies between the localizer and main runs could exists for 3 reasons: 1) reduced power in the main runs compared to localizer runs (i.e. fewer fMRI blocks were included in the contrast), 2) possible differences in the responses to real vs CGI images, and 3) possible differences in the reliabilities of the voxel responses to the different stimulus sets. Moreover, this suggests that reachspaces may not be simply characterized in terms of viewing distance or the size of scale, and there might be other fundamental components missing from our CGI images. Further investigation is necessary to answer these questions.

**Supplementary Figure 6:**
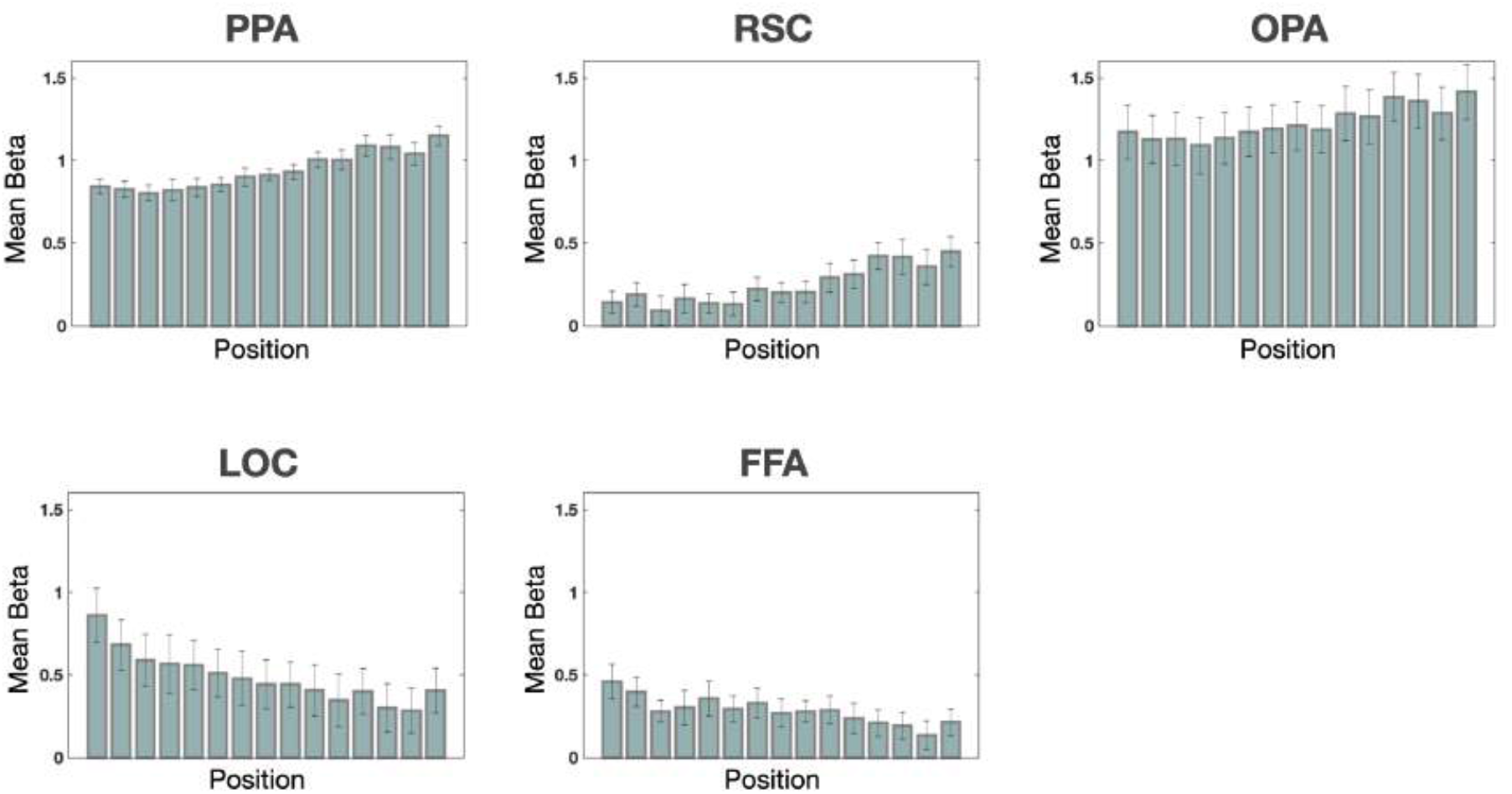
Univariate responses in the ROIs. Each ROI was individually defined using the localizer data. Within each ROI, univariate responses were averaged across the voxels for each Position (i.e., depicted spatial scale), then they were averaged across participants. The group mean betas are plotted from object-view to scene-view along the x-axis. The error bar represents standard error across participants. PPA: parahippocampal place area; RSC: retrosplenial cortex; OPA: occipital place area; LOC: lateral occipital complex; FFA: fusiform face area.

